# How coiled-coil assemblies accommodate multiple aromatic residues

**DOI:** 10.1101/2021.02.01.429152

**Authors:** Guto G. Rhys, William M. Dawson, Joseph L. Beesley, Freddie J. O. Martin, R. Leo Brady, Andrew R. Thomson, Derek N. Woolfson

## Abstract

Rational protein design requires understanding the contribution of each amino acid to a targeted protein fold. For a subset of protein structures, namely the *α*;-helical coiled coils (CCs), knowledge is sufficiently advanced to allow the rational *de novo* design of many structures, including entirely new protein folds. However, current CC design rules center on using aliphatic hydrophobic residues predominantly to drive the folding and assembly of amphipathic *α* helices. The consequences of using aromatic residues—which would be useful for introducing structural probes, and binding and catalytic functionalities—into these interfaces is not understood. There are specific examples of designed CCs containing such aromatic residues, *e.g.*, phenylalanine-rich sequences, and the use of polar aromatic residues to make buried hydrogen-bond networks. However, it is not known generally if sequences rich in tyrosine can form CCs, or what CC assemblies these would lead to. Here we explore tyrosine-rich sequences in a general CC-forming background and resolve new CC structures. In one of these, an antiparallel tetramer, the tyrosine residues are solvent accessible and pack at the interface between the core and the surface. In the other more-complex structure, the residues are buried and form an extended hydrogen-bond network.

---

Coiled coils (CCs) are ubiquitous natural protein-folding and protein-protein-interaction motifs. They have been used extensively as models for protein folding, assembly and design for several decades.^1–6^ As a result, there is a wealth of sequence and structural data on CCs.^7^ This has delivered sequence-to-structure relationships and parametric descriptions for CCs.^8–10^ In turn, these have enabled successful rational and computational CC designs.^11–14^

Despite these extensive studies, surprisingly little has been done to explore the inclusion of aromatic (Ar) residues in the hydrophobic cores of natural or designed CC peptides and proteins.^15–17^ Most likely, this is because early analyses and subsequent experimental studies reveal that different combinations of aliphatic hydrophobic residues—particularly isoleucine (Ile) and leucine (Leu)—in CC interfaces suffice to stabilize and distinguish the predominant oligomeric states of CCs, namely dimers, trimers and tetramers.^18,19^ Indeed, analysis of the current RCSB Protein Data Bank reveals that these two residues account for >40% of all the residues at the traditional hydrophobic core (****a**** and ****d****) sites of all determined CC structures,^7^ which is ≈3 times more than expected by chance. By contrast, aromatic side chains—Phe, Tyr and Trp—found in natural coiled coils (Figure 1A & B) account for <10% of residues at the traditional hydrophobic core positions, approximately as expected by chance.^7^

**Figure 1.**
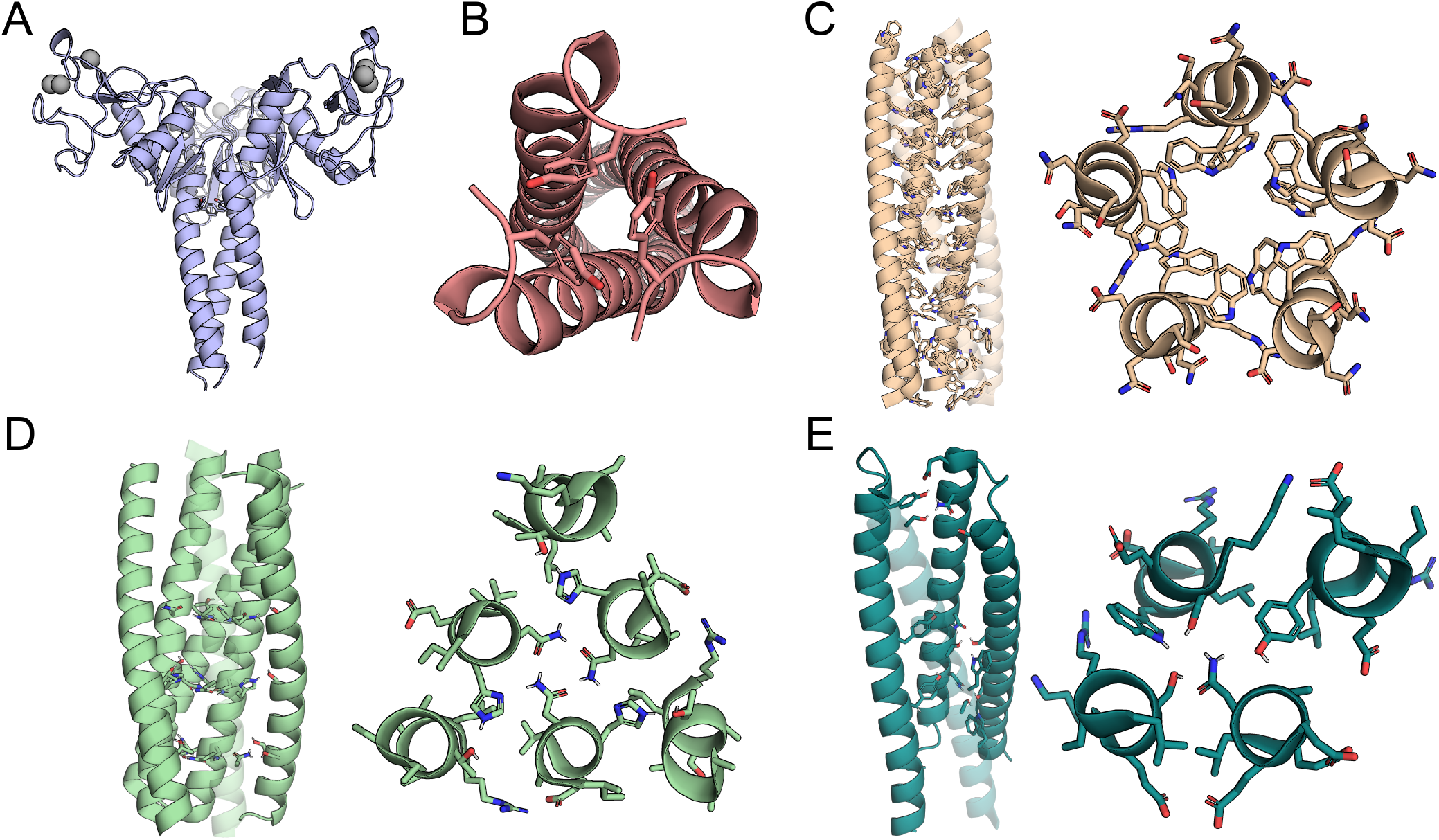
Examples of natural (A&B) and designed (C-E) CCs with buried polar aromatic residues. The structures are (A) the trimeric fragment of human lung surfactant protein D (PDB ID: 1b08); (B) the outer membrane lipoprotein from *E. coli* (PDB ID: 1eq7); (C) an engineered mutant of the outer membrane lipoprotein from *E. coli* (PDB ID: 1t8z); and (D&E) *de novo* designed CCs with buried hydrogen-bond networks, namely (D) pRO-2.5 (PDB ID: 6msr) and (E) DHD131 (PDB ID: 6dkm). In panels C-E some helices are semitransparent to aid visualization of interactions in the core of the helical bundles.

There are some studies of peptides with Ar residues incorporated into CC dimers, trimers and higher-order oligomers.^15–17^ Specific examples of Ar-containing *de novo* and engineered CCs include: engineered pentameric barrels containing buried Trp or Phe residues (Figure 1C),^20,21^ and heterotrimeric CCs with a buried Trp.^22–24^ More recent examples include various parallel and antiparallel helical bundles with consolidated cores that feature Phe residues,^25–27^ and homomeric and heteromeric helix-loop-helix motifs with isolated Tyr, His and Trp residues that contribute to buried hydrogen-bond networks (Figure 1D & E).^14^

In analogous helical systems—such as natural and synthetic DNAs, foldamers, and supramolecular assemblies—π-π stacking and hydrogen-bonding interactions between aromatic moieties contribute to the forces that drive molecular folding, specification and stabilization.^28–30^ Inspired by these studies, and by a gap that we perceived in the CC field, we wondered if normally self-associating CC peptide sequences incorporating multiple Tyr residues at ****a**** or ***d*** sites would assemble to form stable and discrete structures.

Our aim was a test if such inclusions would lead to similar intra- and inter-molecular Ar-Ar interactions observed in the abovementioned analogous systems. Such motifs and interactions in the cores of CC structures could result in desirable properties such as the inclusion of chromophore and fluorophore probes for biophysical studies;^31^ platforms for binding small molecules;^32^ the introduction of sites for post-translational modification;^33^ catalytic centers;^34^ and sub-structures that promote electron transfer.^35^

Recently, largely through rational and computational design, known CC structures have been expanded to larger oligomers of 5 or more helices.^11,12,27,36,37^ Many of these are *α*-helical barrels with central, solvent-accessible channels. We reasoned that Tyr residues might be tolerated better in these barrels and related higher-order structures than in classical dimers to tetramers, as the former have more-expanded helical interfaces and large inter-helical cores or spaces. Therefore, we introduced Tyr and aliphatic residues at the ***a*** and ***d*** sites of a typical *α*-helical barrel, CC-Type2-(L_a_I_d_)_4_, with the sequence repeat LKEIAxA (Table S1).^11^ In addition to the functional reasons given above, we avoided Trp because this has been shown not to be fully tolerated in the cores of some CC assemblies because of its bulk (e.g., PDB ID: 1cos),^38^ and His because burying these residues could lead to potential pH instability of structures.

We combined Tyr residues with the aforementioned aliphatic residues at the core ***a*** and ***d*** positions to generate a series of variants. Whilst many of the resulting peptides were unfolded or aggregated, some formed stable helical assemblies, albeit of lower thermal stability than the parent. X-ray protein crystal structures revealed helical bundles with knobs-into-holes side-chain packing characteristic of CCs, plus buried hydrogen-bond networks. Two designs formed a new type of homomeric CC assembly where symmetry is broken with mixed parallel/antiparallel chain topologies. These studies add to the broader body of the CC design principles and address how multiple aromatic residues—which are not common in natural CCs—might be tolerated in *de novo* CC designs when needed for function.

## EXPERIMENTAL SECTION

### 1.1 Peptide synthesis and purification

#### Peptide synthesis and purification

All Fmoc protected proteinogenic amino acids, DMF and activators were purchased from AGTC Bioproducts, UK. All other solvents were supplied by Fisher Scientific, UK. Chem-Matrix solid supports were supplied by PCAS Biomatrix, Canada. Peptides were synthesized by standard Fmoc SPPS methods on a CEM Liberty Blue automated solid-phase peptide synthesis apparatus with inline UV monitoring. Activation was achieved using DIC/Cl-HOBt. All peptides were produced as the *C-*terminal amide on a Rink amide Chem-Matrix solid support. All peptides were N-terminally acetylated using an excess of acetic anhydride and pyridine in DMF. Cleavage from the solid support was effected with trifluoroacetic acid (TFA) containing 5% triisopropylsilane and 5% water. Following cleavage, the TFA solution was reduced in volume to ≈ 5 ml or lower using a flow of nitrogen. Addition of diethyl ether (≈ 45 ml at 0 °C) gave a precipitate that was recovered via centrifugation and redissolved in a 1:1 solution of acetonitrile in water before freeze-drying, to yield the crude peptide as white or yellow solids. Peptides were purified by reverse-phase HPLC, with a gradient of acetonitrile in water (each containing 0.1% TFA). The stationary phase was a C18 column of dimensions 200 mm by 10 mm. Pure fractions were identified by analytical HPLC and MALDI mass spectrometry, then pooled and freeze-dried.

#### Analytical HPLC for designed sequences

Analytical HPLC traces were obtained using a Jasco 2000 series HPLC system using a Phenomenex ‘Kinetex’ 5 μm particle size, 100 Å pore size, C18 column of dimensions 100 × 4.6 mm. Chromatograms were monitored at 220 and 280 nm wavelengths. The gradient was 20 to 80% acetonitrile in water (each containing 0.1% TFA) over 25 minutes.

#### MALDI-TOF mass spectrometry

MALDI-TOF mass spectra were collected on a Bruker Ultra-Flex MALDI-TOF or ABI 4700 MALDI TOF mass spectrometer operating in positive-ion reflector mode. Peptides were spotted on a ground steel target plate using dihydroxybenzoic acid as the matrix. Masses quoted are for the monoisotopic mass as the singly protonated species. Unless explicitly stated, measured mass is to 0.1% accuracy.

#### Peptide concentration determination

Peptide concentrations were determined by dissolving pure freeze-dried samples in deionized water and measuring the absorption at 280 nm on a Thermo Scientific (Waltham, USA) NanoDrop 2000 UV/Vis spectrometer. The following extinction coefficients were used to determine concentrations: ɛ_Trp 280 nm_ = 5,690 cm^−1^M^−1^ and ɛ_Tyr 280 nm_ = 1,280 cm^−1^M^−1^

### 1.2 Solution-phase biophysics

#### Circular dichroism spectroscopy

Circular dichroism (CD) data were collected on a JASCO J-810 or J-815 spectropolarimeter fitted with a Peltier temperature controller (Jasco UK). Peptide samples were made up as 10 μM or 100 μM solutions in phosphate buffered saline (PBS; 8.2 mM sodium phosphate, 1.8 mM potassium phosphate, 137 mM sodium chloride, 2.7 mM potassium chloride at pH 7.4) or otherwise as stated. CD spectra were recorded in 5 mm or 1 mm path length quartz cuvettes at 20°C. The instruments were set with a scan rate of 100 nm min^−1^, a 1 nm interval, a 1 nm bandwidth and a 1 s response time and scans are an average of 8 scans recorded for the same sample. Blue line plots report 20 °C scans upon cooling the sample from 95 °C. Thermal denaturation data were acquired at 222 nm between 5 °C and 95 °C, with settings as above and a ramping rate of 40 °C hr^−1^ and are a single recording. Baselines recorded using the same buffer, cuvette and parameters were subtracted from each dataset. The spectra were converted from ellipticities (deg) to mean residue ellipticities (MRE, (deg.cm^2^.dmol^−1^.res^−1^)) by normalizing for concentration of peptide bonds and the cell path length using the equation:

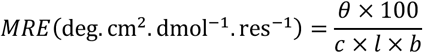

where the variable θ is the measured difference in absorbed circularly polarized light in millidegrees, *c* is the millimolar concentration of the specimen, *l* is the pathlength of the cuvette in cm and *b* is the number of amide bonds in the polypeptide, for which the *N-*terminal acetyl bond was included but not the *C-*terminal amide.

#### Analytical ultracentrifugation

Analytical ultracentrifugation (AUC) sedimentation velocity experiments were conducted at 20 °C in a Beckman Optima XL-A or Beckman Optima XL-I analytical ultracentrifuge using an An-50 or An-60 Ti rotor. Solutions of 310 μl volume were made up in PBS, or otherwise as stated, at 150 μM peptide concentration, and placed in a sedimentation velocity cell with an EPON 2-channel centerpiece and quartz windows. The reference channel was loaded with 325 μl of buffer. The samples were centrifuged at 50k rpm, with absorbance scans taken across a radial range of 5.8 to 7.3 cm at 5 min intervals to a total of 120 scans. Data from a single run were fitted to a continuous c(s) distribution model using Sedfit,^39^ at 95% confidence level. The partial specific volume 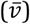 for each of the peptides and the buffer densities and viscosities were calculated using Ultrascan II (http://www.ultrascan.uthscsa.edu). Sednterp (http://rasmb.org/sednterp/) was used to calculate the buffer density and viscosity of solutions of HEPES buffer. Residuals for sedimentation velocity experiments are shown as a bitmap in which the greyscale shade indicates the difference between the fit and raw data. Scans are ordered vertically starting with the first scan at the top. The horizontal axis is the radial range over which the data were fitted.

AUC sedimentation equilibrium experiments were conducted at 20 °C in a Beckman Optima XL-I or XL-A analytical ultracentrifuge using an An-50 Ti or An-60 Ti rotor (Beckmann Coulter). Solutions were made up in PBS, or otherwise explicitly stated, at 150 μM peptide concentration. The experiment was run in triplicate in a 6-channel centerpiece. The samples were centrifuged at speeds in the range 15— 48 krpm and scans at each recorded speed were duplicated. Data were fitted to single, ideal species models using Ultrascan II, comprising a minimum of four speeds. 95% confidence limits were obtained via Monte Carlo analysis of the obtained fits. Peptides that aggregated during the experiment or were deemed unsuitable for analysis as mentioned in the AUC section.

### 1.3 Solid-state analysis

#### Negative-stain transmission electron microscopy

Samples were prepared at 100 μM peptide concentration in PBS (pH 7.4) for 1 h, transferred to glow-discharged 1 nm carbon-coated copper 200 mesh TEM grids (3 μL) and airdried. Grids were stained with 1% uranyl acetate (5 μL, 30 s) and wicked dry with filter paper. Finally, grids were washed with water (5 μL, 30 s) and wicked dry three times and then air-dried. Images were acquired on an FEI Tecnai T12 BioTwin Spirit TEM fitted with an FEI Ceta 4k x 4k CCD camera operating at 120 kV.

#### X-ray crystal structure determination

Freeze-dried peptides were dissolved in deionized water to an approximate concentration of 10 mg ml^−1^ for vapor-diffusion crystallization trials using standard commercial screens (JCSG, Structure Screen 1+2, ProPlex, PACT Premier and Morpheus™) at 19 °C with 0.3 μl of the peptide solution equilibrated with 0.3 μl of the screen solution. To aid with cryoprotection, crystals were soaked in their respective reservoir solutions containing 25% glycerol prior to data collection. X-ray diffraction data were collected at the Diamond Light Source (Didcot, UK) at a wavelength of 0.92 Å on beamlines I03, I04-1 and I03 for CC-Type2-(V_a_Y_d_)_4_-Y3F-W19(BrPhe)-Y24F, CC-Type2-(V_a_Y_d_)_4_-Y3F-W19(BrPhe) and CC-Type2-(Y_a_F_d_)_4_-W19(BrPhe), respectively. Data were processed using the Xia2 3dii autoprocessing pipeline for CC-Type2-(V_a_Y_d_)_4_-Y3F-W19(BrPhe)-Y24F and CC-Type2-(V_a_Y_d_)_4_-Y3F-W19(BrPhe) and the Xia2 Dials pipeline for CC-Type2-(Y_a_F_d_)_4_-W19(BrPhe). The 3dii pipeline uses XDS for point-group selection and integration and XSCALE for scaling and merging of diffraction data.^40^ DIALS pipeline incorporates point-group selection, integration, scaling and merging.^41^ Experimental phasing and structure building was achieved using the Big EP automated pipeline,^42^ which utilizes autoSHARP,^43^ Phenix, AutoSol/AutoBuild^44^ and Crank2.^45^ The datasets were phased with SAD phasing of bromine atoms from 4-bromophenylalanine residues. Final structures were obtained after iterative rounds of model building with COOT^46^ and refinement with Refmac 5.^47^ Solvent-exposed atoms lacking map density were modelled and accounted by high B-factors. All images of crystal structures were generated with PyMOL.^48^

### 1.4 Structure analysis SOCKET analysis

Analysis of knobs-into-hole interaction was performed using the SOCKET server (coiledcoils.chm.bris.ac.uk/socket/). Analysis was conducted on a single biological unit extracted from each pdb file. 7.4 Å was set as the maximum distance of knob residue to a mass averaged position of four hole residues. Perpendicular-like packing was approximated as being 70°< x <110. Acute-like packing was approximated as being 30°< x <70° and 110°< x <150°. Any values outside of these ranges were considered to be parallel-like packing.

#### Analogous structure search

A single biological assembly was generated for each crystal structure. These were submitted to the PDBeFold server (http://www.ebi.ac.uk/msd-srv/ssm/) and CAME TopSearch (https://topsearch.services.came.sbg.ac.at/). Biological Assemblies from PDB release 18/4/2018 was used as the search criteria for CAME TopSearch. The whole PDB archive was searched with PDBeFold without matching connectivity.

#### Tyr-Tyr interaction search

The coiled-coil database, CC+ (coiledcoils.chm.bris.ac.uk/ccplus/search/dynamic_interface) was filtered for non-identical CCs with Tyr at ***a***, ***d***, ***e*** or ***g*** positions. Scripts were written using BioPython.^49^ To measure π–π interactions, centroid/central positions were taken from an average of C*γ* to C*ζ* for each Tyr aromatic ring. Pairwise distances were measured for all Tyr OH–OH atoms and for all π–π in each structure up to a distance of 3.2 Å and 4 Å, respectively. Scripts are available online (https://github.com/woolfson-group/multiple_aromatic_paper_2021).

## RESULTS AND DISCUSSION

### Redesign rationale

We based our target peptide sequences on the computationally designed heptameric *α*-helical barrel, CC-Hept,^11^ systematically named CC-Type2-(L_a_I_d_)_4_. This has proved a productive background for examining sequence-to-structure relationships in CCs.^27^ This 30-residue peptide has four similar seven-residue (heptad) repeats, and *N*-terminal acetylated Gly and *C-*terminal amidated Gly residues known to stabilize α-helical peptides (Table 1).^50,51^ With the repeats labelled ***a→g,*** the ***a, d, e*** and ***g*** positions of CC-Hept—which are occupied by Leu, Ile, Ala and Ala residues, respectively—form a contiguous hydrophobic surface when configured into an *α* helix. This directs and stabilizes helix-helix interactions to form the parallel heptameric barrel with a hydrophobic channel of diameter ≈7 Å. This type of interaction between CC α helices is known as a Type-2 interface.^1,5^ For this study, Tyr was placed at all ***a*** or ***d*** sites of the CC-Type2-(L_a_I_d_)_4_ background, with the other ***a*** or ***d*** position made Ile, Leu, Met, Val, Phe or Tyr to give eleven new sequences. These were named systematically as CC-Type2-(**X**_a_**Z**_d_)_4_, where the **X** and **Z** correspond to the amino acids placed at ***a*** and ***d*** (Table 1). Unless specified, all other positions were maintained from CC-Type2-(L_a_I_d_)_4_. Specifically, the ***e*** and ***g*** positions were Ala to promote formation of large bundles; the solvent-exposed ***b*** and ***c*** sites were Lys and Glu to foster inter-helical salt bridges between α helices; and the exterior ***f*** positions were varied between heptad repeats to maintain solubility (Gln and Lys) and to introduce a chromophore and aid crystallization (Trp).

**Table 1.**
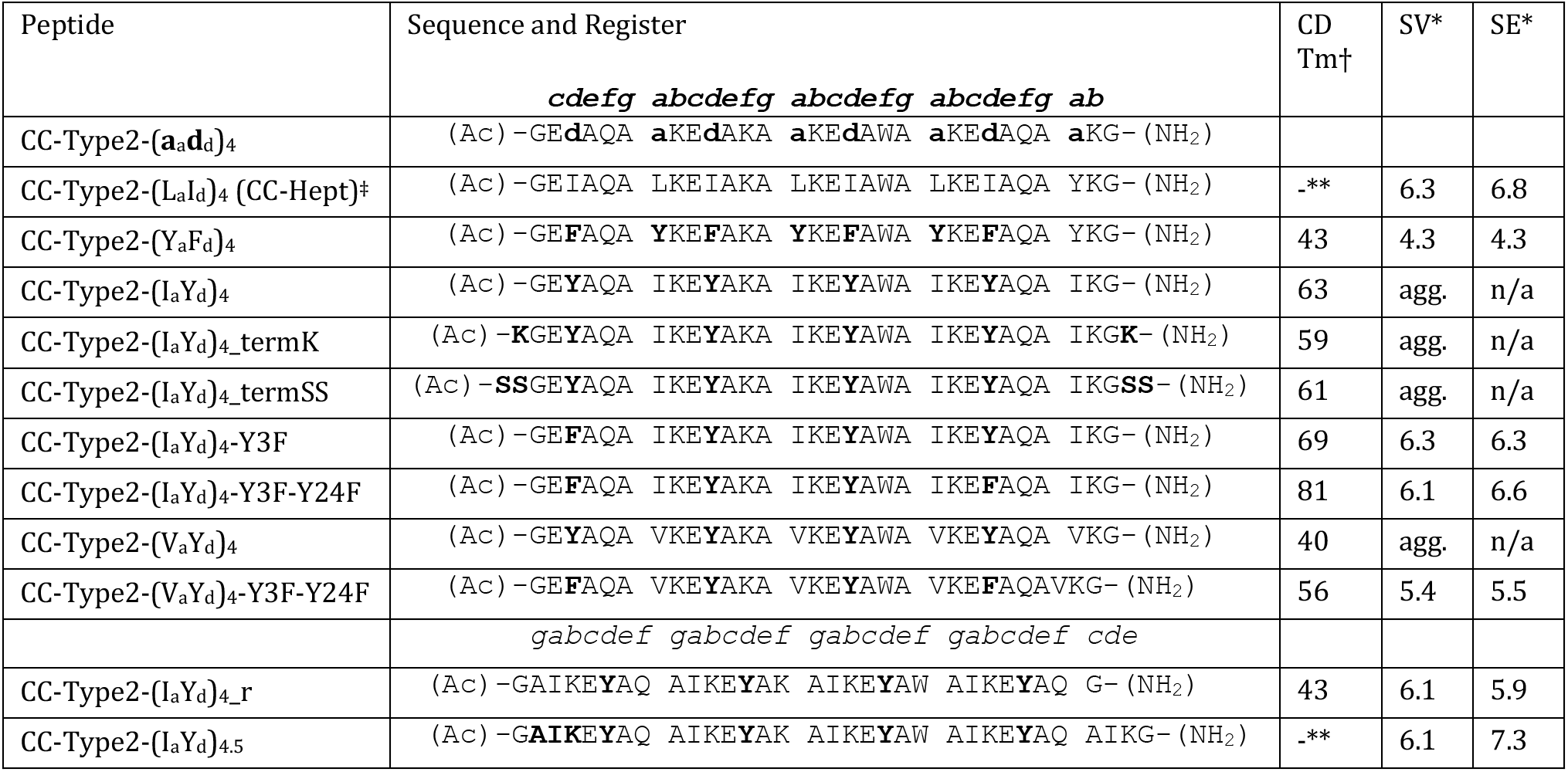
Sequences and biophysical data for folded peptides. A comprehensive table including all unfolded peptides can be found in the supplementary information (Table S1). †Circular dichroism melt-transition temperature. *Sedimentation velocity (SV) and sedimentation equilibrium (SE) analytical ultracentrifugation (mass/monomer mass). **Transition temperature was too high to measure. ‡ Data measured in phosphate-buffered saline solution taken from a previous publication.^11^

Ten sequences were synthesized successfully by solidphase peptide synthesis and purified by reverse-phase HPLC (Table S1) (CC-Type2-(Y_a_M_d_)_4_ was not successfully synthesized). These were characterized by analytical HPLC and MALDI spectrometry (Figures S2.1–2.9).

### Solution-phase characterization

Each peptide was screened for α-helical folding by circular dichroism (CD) spectroscopy at 10 μM and 100 μM peptide concentrations (Figure S3.1). Of the ten peptides, only three were both α helical and showed cooperative unfolding transitions, namely: CC-Type2-(Y_a_F_d_)_4_, CC-Type2-(I_a_Y_d_)_4_ and CC-Type2-(V_a_Y_d_)_4_ (Figure 2A & Figure S3.5, S3.7 & S3.8). Thermal annealing did not significantly improve the α helicity of the other sequences (Figure S3.2-3.4, S3.6, S3.9, S3.10). For CC-Type2-(Y_a_F_d_)_4_, some α helicity was lost after a thermal melt, while the other two were reversible. At 10 μM, CC-Type2-(V_a_Y_d_)_4_, CC-Type2-(Y_a_F_d_)_4_ and CC-Type2-(I_a_Y_d_)_4_ had mid-point thermal unfolding temperatures (TM) of 40 °C, 43 °C and 63 °C, respectively.

**Figure 2.**
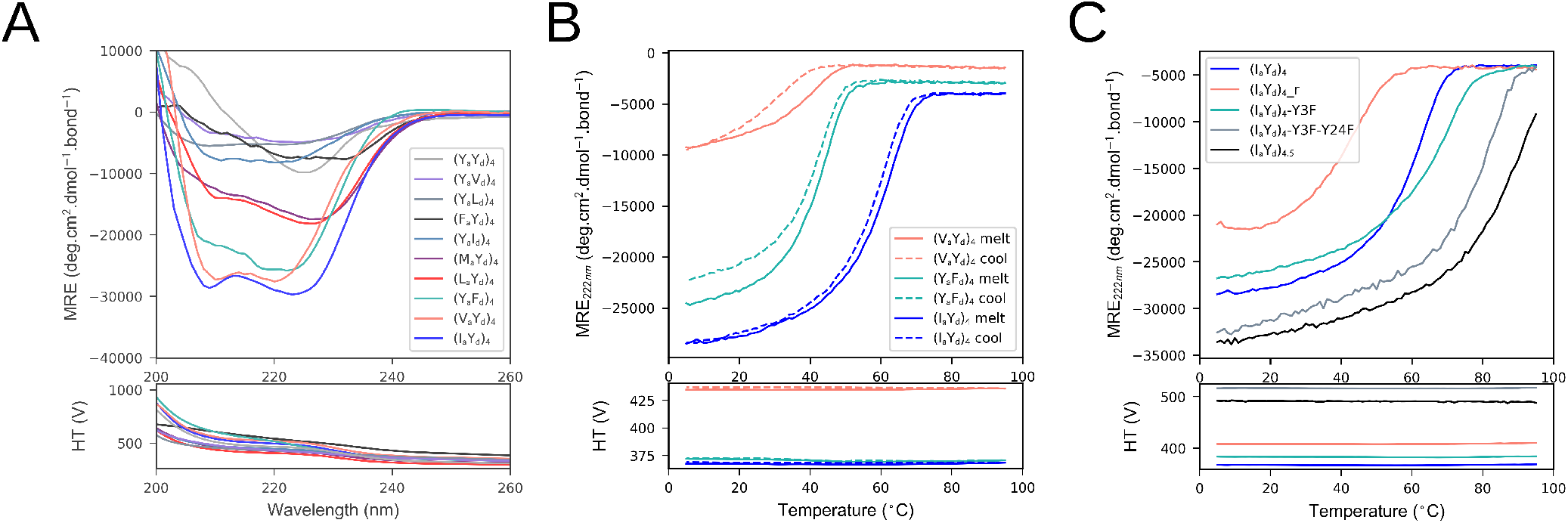
Biophysical characterization of sequences containing Tyr at ***a*** and ***d*** positions. (A) Overlaid circular dichroism spectra for the parent sequences. Spectrum conditions: 100 μM peptide concentration, PBS at pH 7.4. (B) Thermal-denaturation and subsequent cooling profiles for CC-Type2-(I_a_Y_d_)_4_, CC-Type2-(V_a_Y_d_)_4_, CC-Type2-(Y_a_F_d_)_4_, and (C) discrete variants of CC-Type2-(I_a_Y_d_)_4_. (B & C) Thermal-denaturation conditions: 10 μM peptide concentration, PBS at pH 7.4.

To examine polydispersity and oligomeric state, the three folded peptides were subjected to sedimentation-velocity (SV) and sedimentation-equilibrium (SE) experiments using analytical ultracentrifugation (AUC). By both techniques, CC-Type-2-(Y_a_F_d_)_4_ formed a discrete tetramer (Table 1, Figure S4.2). However, CC-Type2-(I_a_Y_d_)_4_ and CC-Type2-(V_a_Y_d_)_4_ were found to aggregate by SV (Figure S4.1). Transmission electron microscopy was used to probe this aggregation further (Figure S5.1 & S5.2), but no defined suprastructures were observed, suggesting that CC-Type2-(I_a_Y_d_)_4_ and CC-Type2-(V_a_Y_d_)_4_ aggregate non-specifically. To test this, we synthesized two variants of CC-Type2-(I_a_Y_d_)_4_ with charged (CC-Type2-(I_a_Y_d_)_4__termK) and polar (CC-Type2-(I_a_Y_d_)_4__termSS) residues at both termini. However, neither prevented aggregation (Figure S4.1).

Another possibility was that, because of the side-chain hydroxyl moieties, the terminal Tyr residues caused fraying of any bundles formed and that this led to aggregation. Fraying is observed in structures of designed CC bundles containing polar residues near their termini and in natural CC structures in the PDB (Figure 1B).^52,53^ To test this, we replaced the most-terminal Tyr residues with Phe in CC-Type2-(I_a_Y_d_)_4_ to give CC-Type2-(I_a_Y_d_)_4_-Y3F & CC-Type2-(I_a_Y_d_)_4_-Y3F-Y24F; and we shifted the register of the sequence (CC-Type2-(I_a_Y_d_)_4__r) or added an extra half heptad (CC-Type2-(I_a_Y_d_)_4.5_) to move those Tyr residues more centrally into the sequences.

Encouragingly, by AUC, all four peptides formed discrete and mostly hexameric species in solution, although a fraction of CC-Type2-(I_a_Y_d_)_4__r aggregated at low centrifugal speeds prior to analysis (Table 1, Figure S4.3-S4.6). The thermal unfolding transitions were close to fully reversible allowing thermal stabilities of the peptides to be measured by CD spectroscopy (Table 1, Figure S3.14-S3.17). Shifting the register had a detrimental effect, lowering the T_M_ by 20°C. Introducing a single Phe gave a modest increase in stability (6 °C), whereas introducing two Phe residues increased the stability by 18 °C. The addition of an extra half-heptad stabilized the assembly such that it did not fully unfold by 95 °C.

To test if a similar strategy could be used to render discrete α-helical bundles for CC-Type2-(V_a_Y_d_)_4_, we synthesized the double Phe mutant (CC-Type2-(V_a_Y_d_)_4_-Y3F-Y24F). Indeed, the thermal stability of this variant improved compared with the parent; and it formed oligomers between pentamer and hexamer by AUC (Table 1. Figures S3.17 and Figure S4.8).

To summarize the solution-phase biophysical data, sequences with multiple Tyr residues at the core ***a*** and ***d*** positions of the CC repeat can form stable and discrete α-helical bundles. For instance, CC-Type2-(Y_a_F_d_)_4_ forms a tetramer. However, and although α helical and co-operatively folded, CC-Type2-(V_a_Y_d_)_4_ and CC-Type2-(I_a_Y_d_)_4_ do not form discrete bundles. This can be remedied by altering ***a*** and ***d*** residues closest to the peptide termini.

### Peptide X-ray crystallography

Crystallization screens were set up for all peptides that formed discrete assemblies in solution. Diffracting crystals were only obtained for CC-Type2-(Y_a_F_d_)_4_ and CC-Type2-(V_a_Y_d_)_4_-Y3F-Y24F. However, the resulting datasets could not be phased. Therefore, variants with the single Trp replaced by 4-Bromo-phenylalanine (BrPhe) were synthesized. The aim was to explore alternative crystal forms, and to provide heavy atoms for single-wavelength anomalous dispersion (SAD) experimental phasing. Gratifyingly, datasets for BrPhe variants of both sequences could be phased.

### An unusual antiparallel CC tetramer

In agreement with solution-phase data for CC-Type2-(Y_a_F_d_)_4_, its X-ray crystal structure revealed a tetramer (Figure 3). More specifically, CC-Type2-(Y_a_F_d_)_4_-W19(BrPhe) formed an oblate D_2_ symmetric antiparallel tetramer. Like our previously *de novo* designed antiparallel tetramer with aliphatic residues at ***a*** and ***d***, apCC-Tet,^52^ the oblate structure has two ‘narrow interfaces’ (Figure 3A) and two ‘wide interfaces’ (Figure 3B). The narrow interface is made by the Ala residues at ***g***, the Phe residues form the core, and the Tyr residues reach across the wide interfaces (Figure 3D & E).

**Figure 3.**
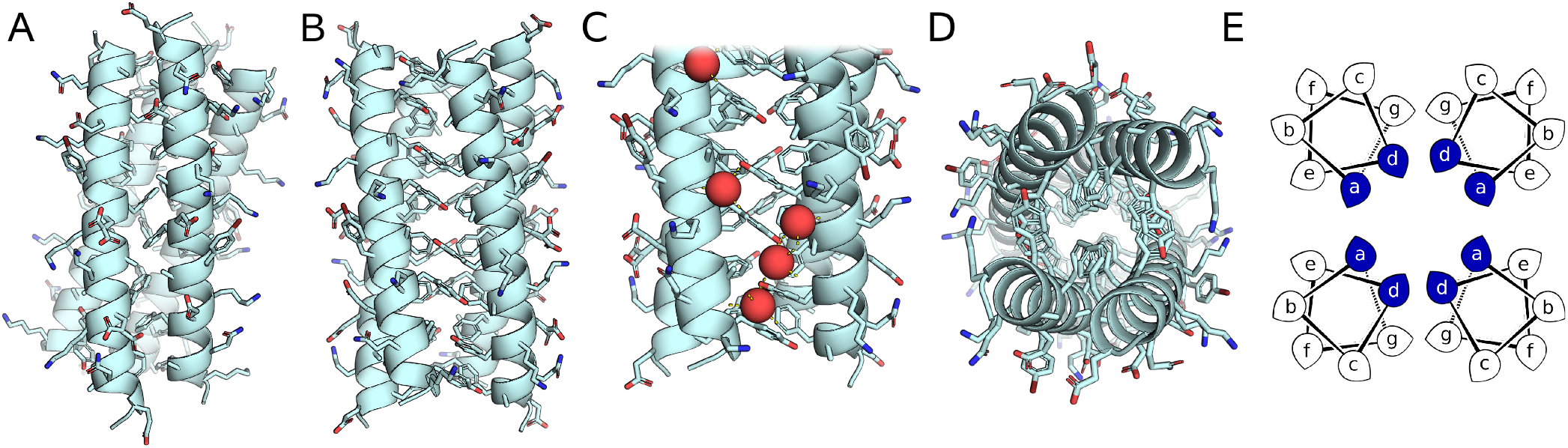
X-ray crystal structure of CC-Type2-(Y_a_F_d_)_4_-W19(4BrPhe) viewed from the short interface (A), the long interface (B&C), and along the superhelical axis (D). (C) Surface waters (red spheres) interacting with tyrosine side chains. (E) Helical-wheel diagram with knob residues colored blue.

To confirm the structure as a CC, we analyzed it for knobs-into-holes (KIH) interactions using SOCKET.^54^ This revealed interactions dominated by the aromatic residues. Apart from the *N-*terminal Phe and the *C-*terminal Tyr, which cannot interact with complete sets of hole residues, all of the aromatic residues acted as knobs. Surprisingly, the aforementioned Ala residues at the ***g*** positions do not act as classical knob residues according to SOCKET. Visual inspection revealed that these Ala residues in the narrow interface pack in ***g***-to-***g*** knob-to-knob interactions (Figure 3A). This contrasts with apCC-Tet, where the analogous Ala at ***e*** interdigitate to form a tight interface.^52^ The looser Ala interface of CC-Type2-(Y_a_F_d_)_4_-W19(BrPhe) can be attributed to the offset of the α helices required to accommodate the aromatic residues in the wide interface (Figure 3B).

Indeed, the phenolic side chains interact in a specific manner. Each Tyr pairs antiparallel with another from an adjacent α helix. The average Tyr-Tyr aromatic-centroid distances (Supplementary Information 1.4) are 3.8 Å (± 0.1 Å) and 3.9 Å (± 0.2 Å), which are at the upper-limit of recognized distances for π-π stacking.^55,56^ Also, the hydroxyl groups interact with water molecules at the wide interface to form a hydrogen-bond network (Figure 3C).

### A new complex CC with polar core

By contrast, the X-ray crystal structure of the triple mutant CC-Type2-(V_a_Y_d_)_4_-Y3F-W19(BrPhe)-Y24F revealed a low-symmetry hexameric bundle with five parallel α helices and a single antiparallel α helix (Figure 4A). As described below, the helices are offset from each other along the superhelical axis. This appears to facilitate formation of a core with an extensive buried hydrogen-bonded network, which involves two water molecules and a polyethylene glycol molecule (Figure 4B).

**Figure 4.**
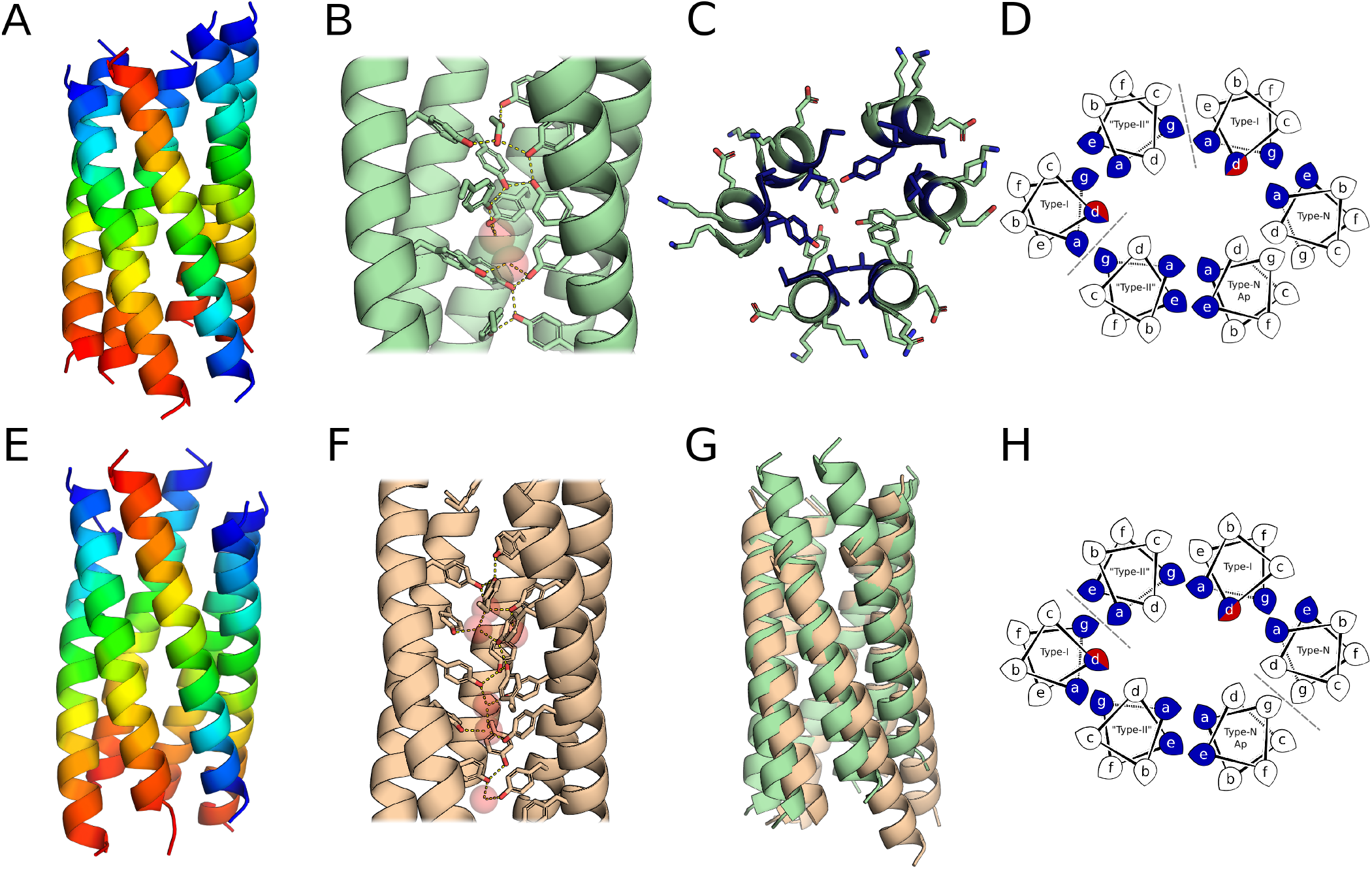
Crystal structures and analysis of CC-Type2-(V_a_Y_d_)_4_ variants with buried tyrosine hydrogen-bond networks. Crystal structures of the triple mutant CC-Type2-(V_a_Y_d_)_4_-Y3F-Y24F-W19(BrPhe) (A-C), and the double mutant CC-Type2-(V_a_Y_d_)_4_-Y3F-W19(BrPhe) (E-F). (A & E) Chains colored in a rainbow spectrum from the blue *N* termini to the red *C* termini. (B & F) Buried hydrogen-bond networks found within the mutants. (C) Two α-helical turns of the triple mutant highlighting knob residues in blue. (D & H) Helical-wheel diagrams of the triple (D) and double mutants (H). Blue and red residues signify the Tyr that form knob residues in both neighboring α helices. Ap denotes the antiparallel α helix and the dashed line indicate interfaces where the α helices are offset by a α helical turn. (G) Structural alignment of both structures ignoring 20 residues at the termini. Blue residues signify knob residues.

SOCKET identified a variety of CC interactions, including Type-N, Type-I and Type-II-like interfaces (Figure 4C & D). One antiparallel interface lacks KIH interactions completely and is preserved solely by the interdigitation of aromatic residues. As in the tetrameric structure of CC-Type2-(Y_a_F_d_)_4_-W19(BrPhe), in this interface the Ala residues at ***g*** are offset and do not interdigitate. The two offset parallel interfaces (Figure 4D, dashed lines) allow the Tyr residues to form knobs to the neighboring α helices. The packing angles of knobs varied from 45° to 135°, which range from what are often termed ‘acute’ through ‘perpendicular’ packing,^54^ is broader than usually observed for any one type of KIH interaction.

Thus, the structure of CC-Type2-(V_a_Y_d_)_4_-Y3F-W19(BrPhe)-Y24F cannot be considered a classical (highly symmetric) CC in any sense. We posit that the complexity of the structure arises from the need to satisfy the hydrogen-bonding propensity of the Tyr residues, and to form the buried network observed. For this to occur, the two parallel interfaces are offset by a single helical turn (half of a CC heptad repeat), and one helix associates with the assembly in an anti-parallel fashion. This accommodation of buried tyrosine side chains is dominated by the drive to form hydrogen bonds rather than by optimizing KIH packing; although such interactions can and are made by the Val residues at ***a***. Finally, it is interesting to compare this structure and the tetramer of CC-Type2-(Y_a_F_d_)_4_-W19(BrPhe), which has Tyr at ***a*** and rather than ***d***. It appears that the Tyr at ***a*** has an ‘escape route’ in the tetramer that allows the hydroxy moieties to interact with bulk solvent, but that this is not accessible with Tyr at ***d***.

### Subtle variations on a theme

With the successful structure determination of the triple variant CC-Type2-(V_a_Y_d_)_4_-Y3F-Y24F-W19(BrPhe), we sought to address issues that we had encountered earlier in the study regarding bundle fraying and aggregation. We made a variant with a single Y→ F change, namely CC-Type2-(V_a_Y_d_)_4_-Y3F-W19(BrPhe) (Table S1), and solved its X-ray crystal structure (Figures 4E-H). At first glance, the structure is similar to the triple variant: it is a collapsed hexameric bundle with a single antiparallel chain; the core has an extended hydrogen-bonded network with buried water molecules; and SOCKET analysis identifies the same knob residues (Figures 4D & H).

However, and despite these similarities, good structural alignment of the single and double Y→ F variants was only possible by omitting twenty residues (Q score, 0.76; RMSD over C*α* atoms, 0.5 Å). Closer examination revealed the interfaces with the offset α helices to be located between different α helices in the two structures (Figures 4D & H). The effect of this is to shift two α helices an entire heptad (two α-helical turns) between the two structures (Figure 4G). This can be understood by comparing the two hydrogenbond networks (Figure 4B & F). While the centers of the networks are largely preserved, the different offsets allow different terminal residues to become exposed. In the triple variant, CC-Type2-(V_a_Y_d_)_4_-Y3F-Y24F-W19(BrPhe), the termini are more flush reducing the solvent exposure of two near-terminal Phe side chains, whereas CC-Type2-(V_a_Y_d_)_4_-Y3F-W19(BrPhe), with one of these Phe replaced by Tyr in each chain, the termini are more ragged allowing some of the additional Tyr residues to become more solvent exposed (Figure 4E & F).

### Similar structures in the RCSB PDB

Intrigued by the complexity of these Tyr-containing structures, we searched PDBeFold and CAME TopSearch for similar protein folds.^57,58^

Unsurprisingly, searches using the tetrameric CC-Type2-(Y_a_F_d_)_4_-W19(4BrPhe) returned a plethora of structures. These included natural and *de novo* designed tetrameric (PDB IDs: 2b1f, 4pxu, 6c52, and 6q5s), dimeric (PDB ID: 3ja6, 4d02, 5szg, and 6dlm), and single-chain (PDB ID: 5g05, 5tgw & 5tgy) four-helix bundles. Despite this abundance of hits, none of the highest-ranking structures contained Tyr buried at interface positions.

Searches using the two structures of the CC-Type2-(V_a_Y_d_)_4_ variants both returned similar hits to each other. The highest-ranking similar structures were dominated by all-parallel CC α-helical barrels, including: natural pentameric barrels, *e.g.* cartilage oligomerization matrix protein (PDB ID: 3v2r); designed hexameric barrels, *e.g.* CC-Hex2 (PDB ID: 4pn9); and designed blunt-ended and engineered slipped heptameric barrels, *e.g.* CC-Hept and GCN4-pAA (PDB IDs: 4pna & 2hy6). Therefore, the CC-Type2-(V_a_Y_d_)_4_ variants have low structural similarity to all protein folds found in the RCSB PDB,^59^and are further evidence of these being in hitherto uncharted territory of protein folds explored by nature.^60,61^

Finally, we assessed if local Tyr–Tyr interactions of the type found in our new structures exist more widely in CC structures. To do this, we searched 2,662 CC assignments from the CC+ database (January 2020),^7^ and found 1,861 assignments (from 1,453 structures) that have Tyr at either ***a***, ***d***, ***e*** or ***g*** positions. In this subset, we searched for potential Tyr– Tyr contacts within the CC regions.

Visual inspection of the hits identified five examples of hydroxyl–hydroxyl distances within 3.2 Å, and another five examples of π–π interactions within 4 Å (Table S8.1, Figure S7.1 & S7.2). Thus, with just ten observations from all 2,662 CC assignments (Figure S7.1), it appears that Tyr-Tyr interactions are rare in CC structures resolved to date. It is possible that such interactions are more probable in higher-order oligomers, like those described herein, as these structures provide more opportunities and more buried surfaces to foster them. Indeed, our CC+ dataset was dominated by dimers (91%), and four of the π–π interactions identified cause significant distortion at dimeric interfaces (Figure S7.2). Turning this around, it may be possible to direct and specify CC assemblies, and particularly higher-order assemblies, by employing aromatic residues and contacts between them in some way.

## CONCLUSION

Designing coiled-coil (CC) peptides that are rich in aromatic side chains is challenging. Here we demonstrate that stable, discrete CC assemblies can be formed from sequences rich in Tyr at buried or partially buried positions. Through this, we observe two modalities for accommodating these side chains. In the first—which is in an antiparallel tetramer with Tyr at the ***a*** sites of a CC sequence repeat—Tyr residues pair at the interface between the hydrophobic core and the CC surface to form π-π interactions and leaving their hydroxyl groups exposed to solvent. In the second, with Tyr at the ***d*** sites, the peptides form complex hexameric CCs with buried Tyr side chains. In these, the hydroxyl groups are accommodated in extended hydrogenbond networks. These Tyr-Tyr interactions add to the palette of interactions that can be used for the *de novo* design of proteins and protein-protein interactions.

The inability to crystallize many of the sequences that we explored, suggests that these may form heterogenous mixtures of α-helical bundles in solution. This highlights the complexity of the CC energetic landscape, where multiple near-isoenergetic structures likely exist.^52^ Therefore, it is essential that we develop new design principles and rules in order to design aromatic-rich CC sequences.

One such principle appears to be the avoidance of placing residues like Tyr where alternative states can be observed. For example, we find that ‘capping the cores’ of Tyr-containing α-helical bundles with Phe rather than Tyr made the peptides easier to handle and more discrete. We attribute this to Phe consolidating and capping the core, as compared with Tyr, which favors interactions of its hydroxyl group with solvent.

From previous studies, we know that some CC sequences based on aliphatic residues preferentially form α-helical barrels with solvent-accessible channels.^27^ Therefore, it is somewhat surprising that we did not observe such assemblies with the Tyr-based sequences. It is possible that sequences containing multiple Tyr prefer to form structures with consolidated cores because of increased aromatic-based hydrophobic interactions. Alternatively, barrel states might be accessible to Tyr-rich sequences, but that other residues specify against it. In this study, the three structures contain Phe at some of the ***a*** or ***d*** positions, which has previously been shown in most cases to form structures with consolidated cores.^25–27^ If this is true generally, designing Tyr-containing sequences without Phe might lead to barrel structures.

Finally, macromolecular structures that are rich in constrained and clustered electron-rich aromatic groups can have interesting optical and electrochemical properties.^62^ Therefore, beyond developing new design rules, structures of the type reported here could form the basis of protein designs, nanomaterials, and sensors with new and useful properties.

## ASSOCIATED CONTENT

### Supporting Information

The Supporting Information is available free of charge on the ACS Publications website.

Supporting Figures and Tables (PDF)

### Crystal-Structure Files

The crystal structures are all available from the Protein Data Bank using the following accession codes: 7boa for CC-Type2-(Y_a_F_d_)_4_-W19(BrPhe), 7bo9 for CC-Type2-(V_a_Y_d_)_4_-Y3F-W19(BrPhe), and 7bo8 for CC-Type2-(V_a_Y_d_)_4_-Y3F-W19(BrPhe)-Y24F.

## AUTHOR INFORMATION

### Author Contributions

G.G.R., A.R.T. and D.N.W. conceived the project and designed the experiments. G.G.R., and W.M.D. synthesized the peptides and conducted the solution-phase biophysics. J.L.B. collected and analyzed transmission electronic microscopy data. G.G.R., F.M. and R.L.B. determined the peptide X-ray crystal structures. G.G.R. and D.N.W. wrote the paper. All authors have read and contributed to the preparation of the manuscript.

### Notes

The authors declare no competing financial interest.

## ACKNOWLEDGMENT

We thank the University of Bristol School of Chemistry Mass Spectrometry Facility for access to the EPSRC-funded Bruker Ultraflex MALDI-TOF/TOF instrument (EP/K03927X/1). We are grateful to BrisSynBio for access to the BBSRC-funded peptide synthesizers (BB/L01386X/1). We would like to thank Diamond Light Source for access to beamlines I03 and I04-1 (Proposal 12342), and for the support from the macromolecular crystallography staff.

## ABBREVIATIONS

CC: coiled coil
KIH: knobs-into-holes.

---

TOC

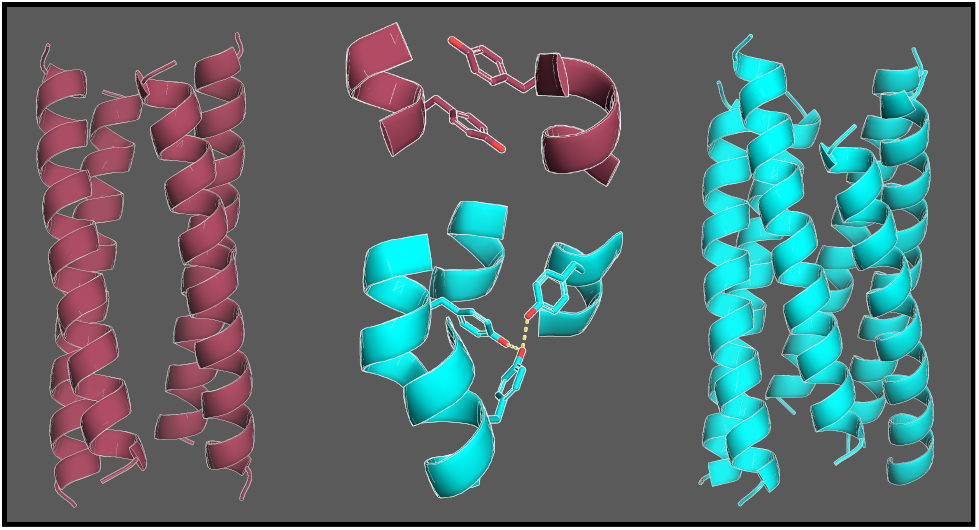

