## Supplementary Information for "How coiled-coil assemblies accommodate multiple aromatic residues"

### Table of Contents

**Table of Contents** **1**

**1 Sequence table** **2**

**2 Matrix-assisted laser desorption/ionisation - time of flight (MALDI-TOF) and Analytical high-pressure liquid chromatography (HPLC)** **3**

**3 Circular dichroism (CD)** **10**

**4 Analytical ultracentrifuge (AUC)** **21**

**5 Negative-stain transmission electron microscope (TEM)** **26**

**6 Crystallography** **27**

**7 Bioinformatics** **29**

### 1 Sequence table

| CC-Type2- | Sequence |
|---|---|
| Initial Screen | *cdefgab cdefgab cdefgab cdefgab* |
| $(Y_aI_d)_4$ | Ac-G EIAQAYK EIAKAYK EIAWAYK EIAQAYK G-$NH_2$ |
| $(Y_aL_d)_4$ | Ac-G ELAQAYK ELAKAYK ELAWAYK ELAQAYK G-$NH_2$ |
| $(Y_aV_d)_4$ | Ac-G EVAQAYK EVAKAYK EVAWAYK EVAQAYK G-$NH_2$ |
| $(Y_aF_d)_4$ | Ac-G EFAQAYK EFAKAYK EFAWAYK EFAQAYK G-$NH_2$ |
| $(Y_aM_d)_4$ | Ac-G EMAQAYK EMAKAYK EMAWAYK EMAQAYK G-$NH_2$ |
| $(L_aY_d)_4$ | Ac-G EYAQALK EYAKALK EYAWALK EYAQALK G-$NH_2$ |
| $(I_aY_d)_4$ | Ac-G EYAQAIK EYAKAIK EYAWAIK EYAQAIK G-$NH_2$ |
| $(V_aY_d)_4$ | Ac-G EYAQAVK EYAKAVK EYAWAVK EYAQAVK G-$NH_2$ |
| $(M_aY_d)_4$ | Ac-G EYAQAMK EYAKAMK EYAWAMK EYAQAMK G-$NH_2$ |
| $(F_aY_d)_4$ | Ac-G EYAQAFK EYAKAFK EYAWAFK EYAQAFK G-$NH_2$ |
| $(Y_aY_d)_4$ | Ac-G EYAQAYK EYAKAYK EYAWAYK EYAQAYK G-$NH_2$ |
| $(Y_aF_d)_4$-W19(BrPhe) | Ac-G EFAQAYK EFAKAYK EFA**X**AYK EFAQAYK G-$NH_2$ |
| $I_aY_d$ optimisation | |
| $(I_aY_d)_4$_termK | Ac-**K**G EYAQAIK EYAKAIK EYAWAIK EYAQAIK G**K**-$NH_2$ |
| $(I_aY_d)_4$_termSS | Ac-**SS**G EYAQAIK EYAKAIK EYAWAIK EYAQAIK G**SS**-$NH_2$ |
| $(I_aY_d)_4$-Y3F | Ac-G E**F**AQAIK EYAKAIK EYAWAIK EYAQAIK G-$NH_2$ |
| $(I_aY_d)_4$-Y3F-Y24F | Ac-G E**F**AQAIK EYAKAIK EYAWAIK E**F**AQAIK G-$NH_2$ |
| | *gabcdef gabcdef gabcdef gabcdef gab* |
| $(I_aY)_4$_r | Ac-G AIKEYAQ AIKEYAK AIKEYAW AIKEYAQ G-$NH_2$ |
| $(I_aY)_{4.5}$ | Ac-G **AIK**EYAQ AIKEYAK AIKEYAW AIKEYAQ AIK G-$NH_2$ |
| $V_aY_d$ optimisation | *cdefgab cdefgab cdefgab cdefgab* |
| $(V_aY_d)_4$-Y3F-Y24F | Ac-G E**F**AQAVK EYAKAVK EYAWAVK E**F**AQAVK G-$NH_2$ |
| $(V_aY_d)_4$-Y3F-W19(BrPhe)-Y24F | Ac-G E**F**AQAVK EYAKAVK EYA**X**AVK E**F**AQAVK G-$NH_2$ |
| $(V_aY_d)_4$-Y3F-W19(BrPhe) | Ac-G E**F**AQAVK EYAKAVK EYA**X**AVK EYAQAVK G-$NH_2$ |

**Table S 1.1** Peptide sequences discussed in the article with their corresponding name. Mutations and additions are shown in bold. X represents 4-bromo-phenylalanine.

### 2 Matrix-assisted laser desorption/ionisation - time of flight (MALDI-TOF) and Analytical high-pressure liquid chromatography (HPLC)

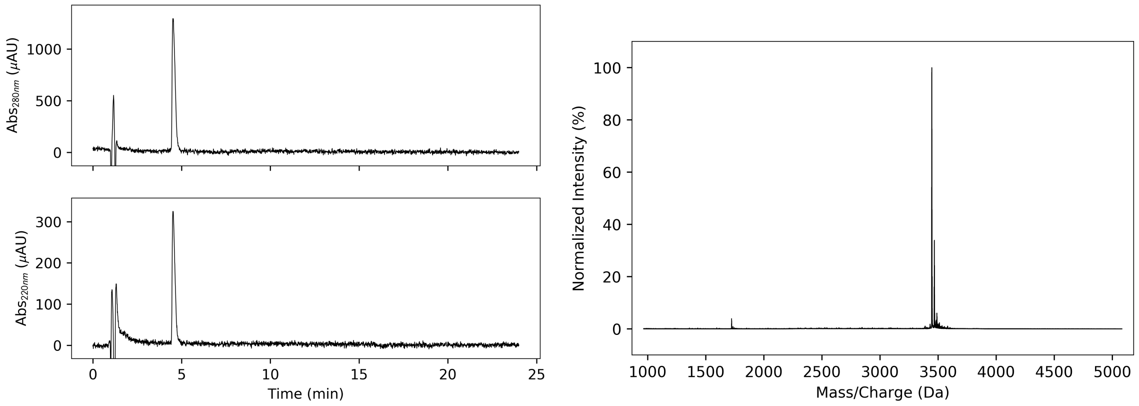


**Figure S 2.1 CC-Type2-($Y_aI_d$)$_4$** - HPLC traces (left, 220 and 280 nm) and MALDI-TOF MS (right). Calculated mass = 3445.8 Da, observed mass = 3447 Da.

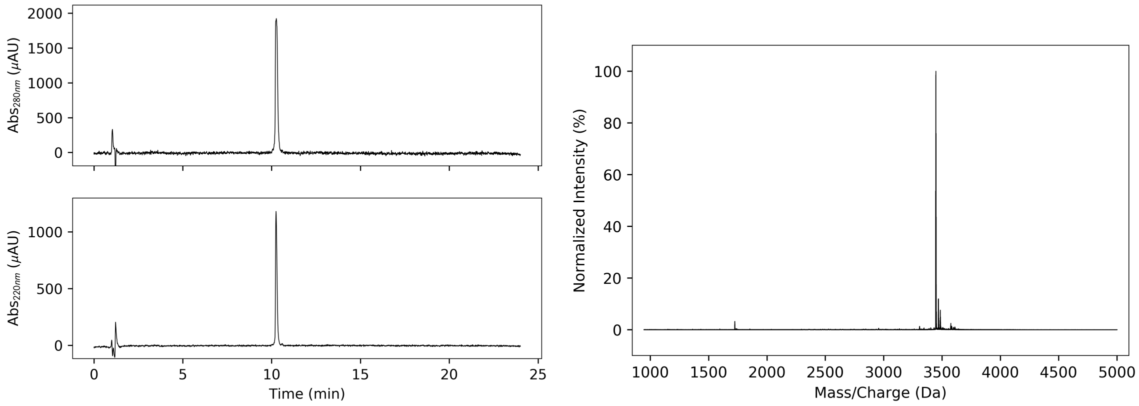


**Figure S 2.2 CC-Type2-($Y_aL_d$)$_4$** - HPLC traces (left, 220 and 280 nm) and MALDI-TOF MS (right). Calculated mass = 3445.8 Da, observed mass = 3446 Da.

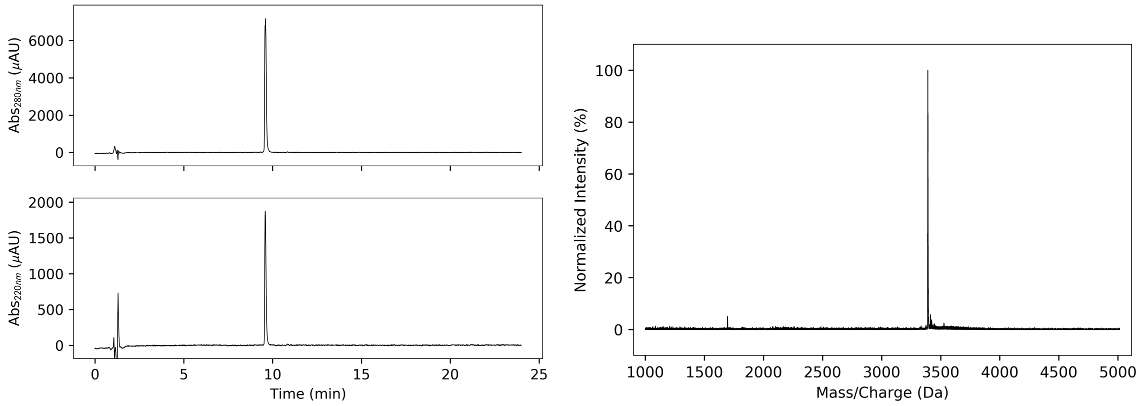


**Figure S 2.3 CC-Type2-$(Y_aV_d)_4$** - HPLC traces (left, 220 and 280 nm) and MALDI-TOF MS (right). Calculated mass = 3389.8 Da, observed mass = 3390 Da.

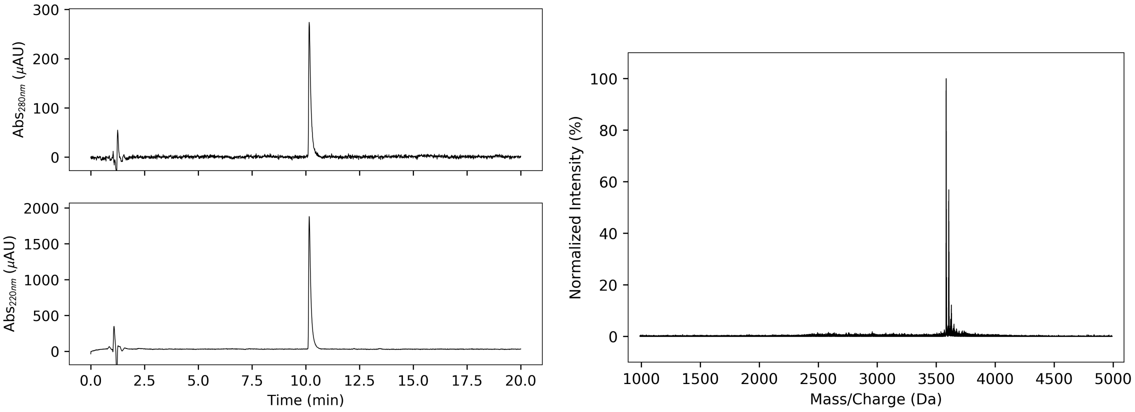


**Figure S 2.4 CC-Type2-$(Y_aF_d)_4$** - HPLC traces (left, 220 and 280 nm) and MALDI-TOF MS (right). Calculated mass = 3580.7 Da, observed mass = 3583 Da.

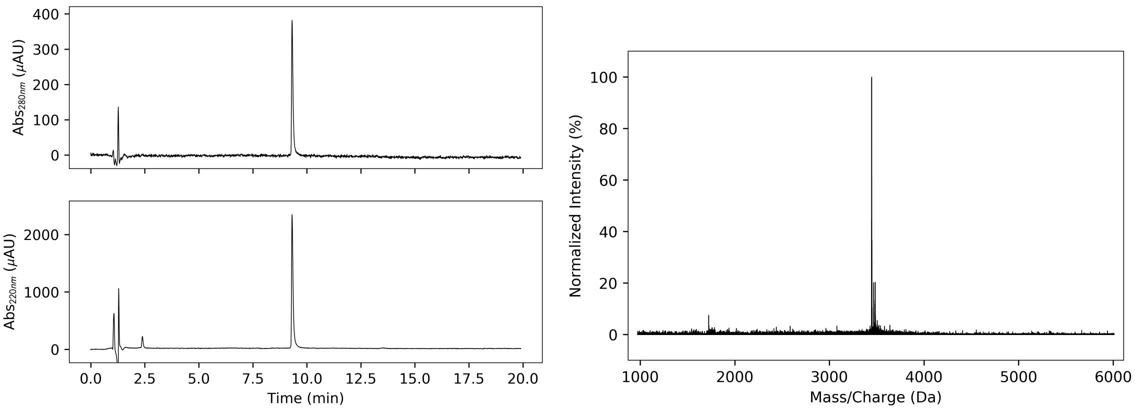


**Figure S 2.5 CC-Type2-$(L_aY_d)_4$** - HPLC traces (left, 220 and 280 nm) and MALDI-TOF MS (right). Calculated mass = 3444.8 Da, observed mass = 3445 Da.

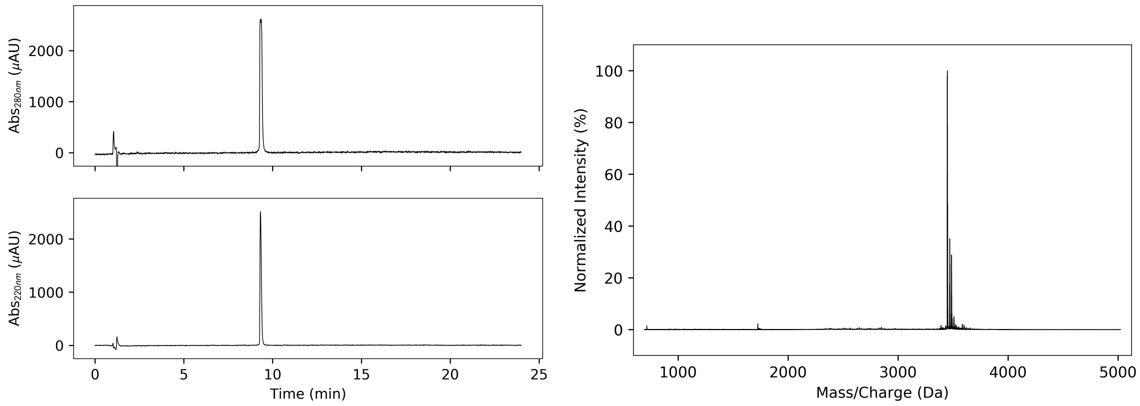


**Figure S 2.6 CC-Type2-($I_aY_d$)$_4$** - HPLC traces (left, 220 and 280 nm) and MALDI-TOF MS (right). Calculated mass = 3445.8 Da, observed mass = 3446 Da.

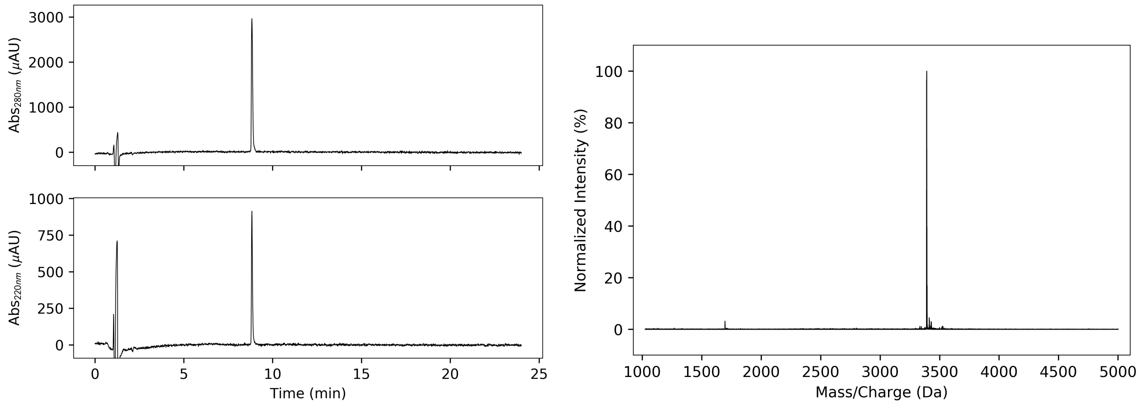


**Figure S 2.7 CC-Type2-($V_aY_d$)$_4$** - HPLC traces (left, 220 and 280 nm) and MALDI-TOF MS (right). Calculated mass = 3389.8 Da, observed mass = 3390 Da.

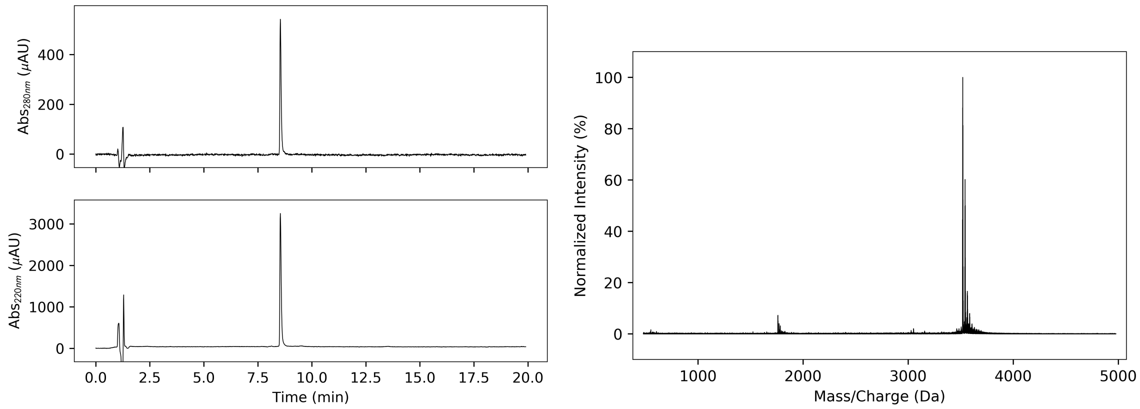


**Figure S 2.8 CC-Type2-($M_aY_d$)$_4$** - HPLC traces (left, 220 and 280 nm) and MALDI-TOF MS (right). Calculated mass = 3516.6 Da, observed mass = 3518 Da.

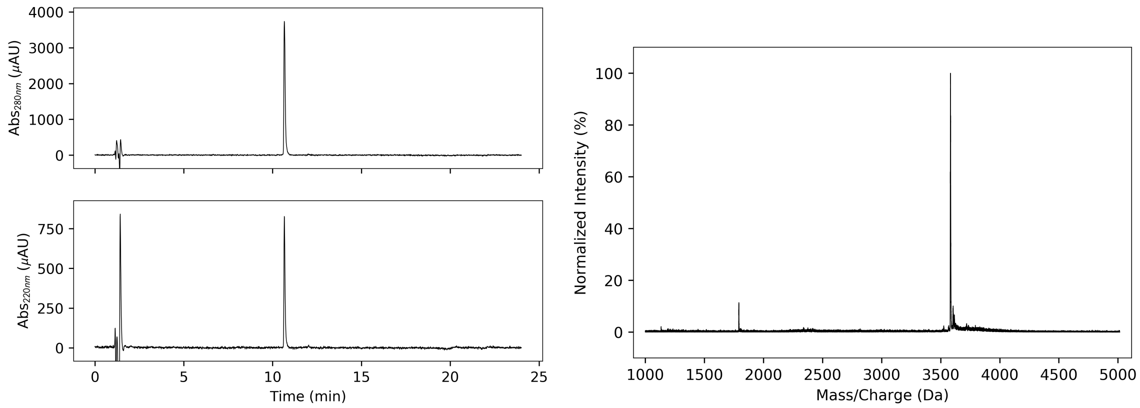


**Figure S 2.9 CC-Type2-($F_aY_d$)$_4$** - HPLC traces (left, 220 and 280 nm) and MALDI-TOF MS (right). Calculated mass = 3581.8 Da, observed mass = 3581 Da.

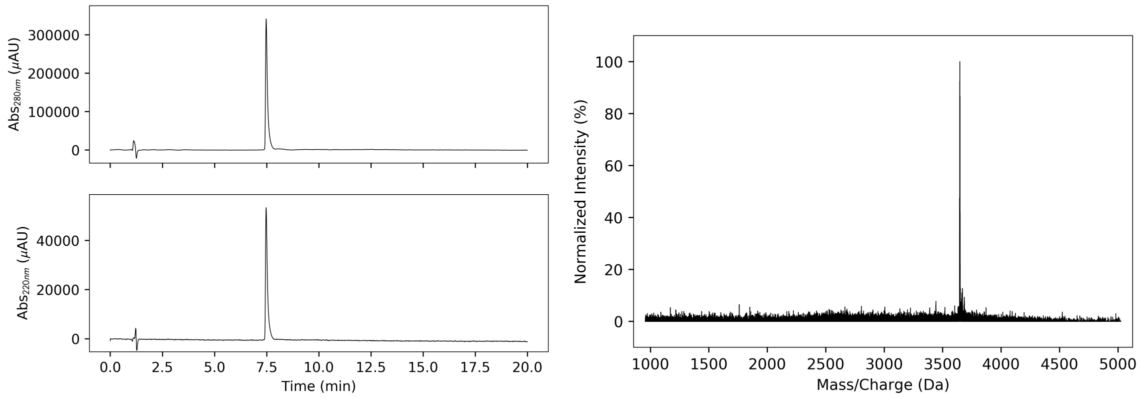


**Figure S 2.10 CC-Type2-($Y_aY_d$)$_4$** - HPLC traces (left, 220 and 280 nm) and MALDI-TOF MS (right). Calculated mass = 3645.7 Da, observed mass = 3646 Da.

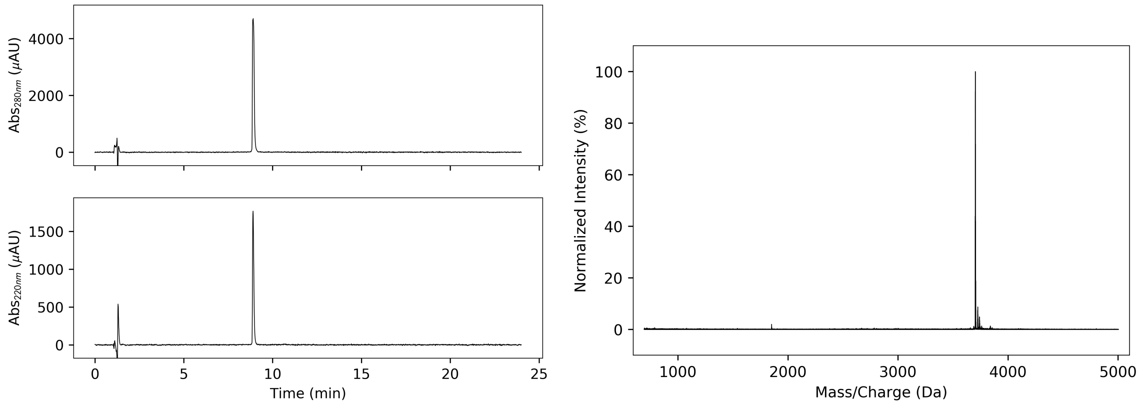


**Figure S 2.11 CC-Type2-($I_aY_d$)$_4$_ termK** - HPLC traces (left, 220 and 280 nm) and MALDI-TOF MS (right). Calculated mass = 3702.0 Da, observed mass = 3702 Da.

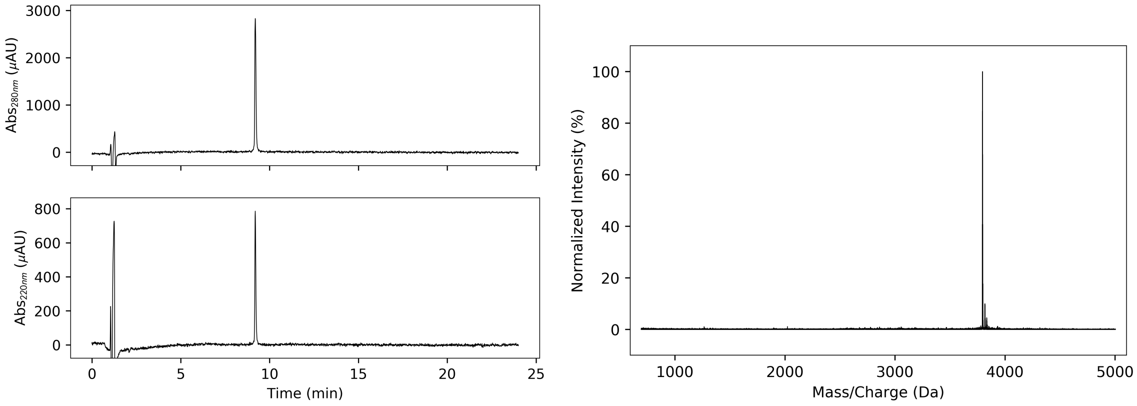


**Figure S 2.12 CC-Type2-$(\mathbf{I_aY_d})_4$_ termSS** - HPLC traces (left, 220 and 280 nm) and MALDI-TOF MS (right). Calculated mass = 3794.0 Da, observed mass = 3794 Da.

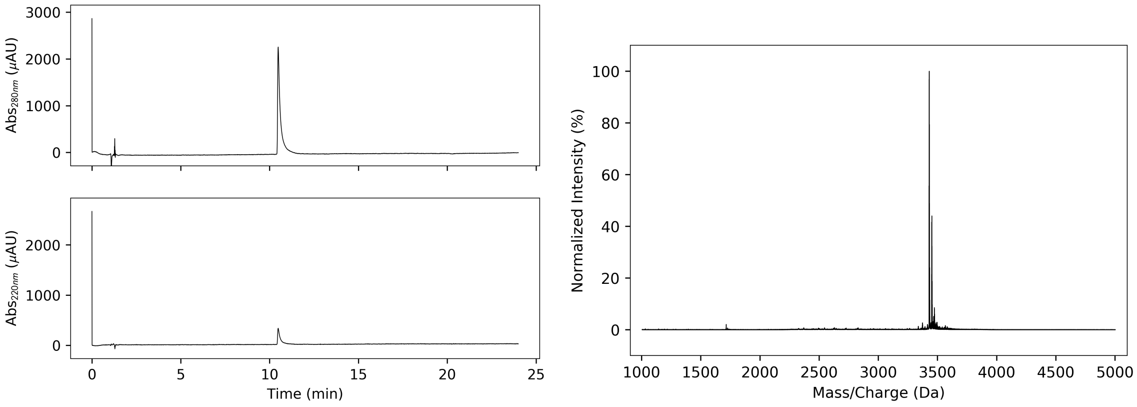


**Figure S 2.13 CC-Type2-$(\mathbf{I_aY_d})_4$-Y3F** - HPLC traces (left, 220 and 280 nm) and MALDI-TOF MS (right). Calculated mass = 3429.8 Da, observed mass = 3431 Da.

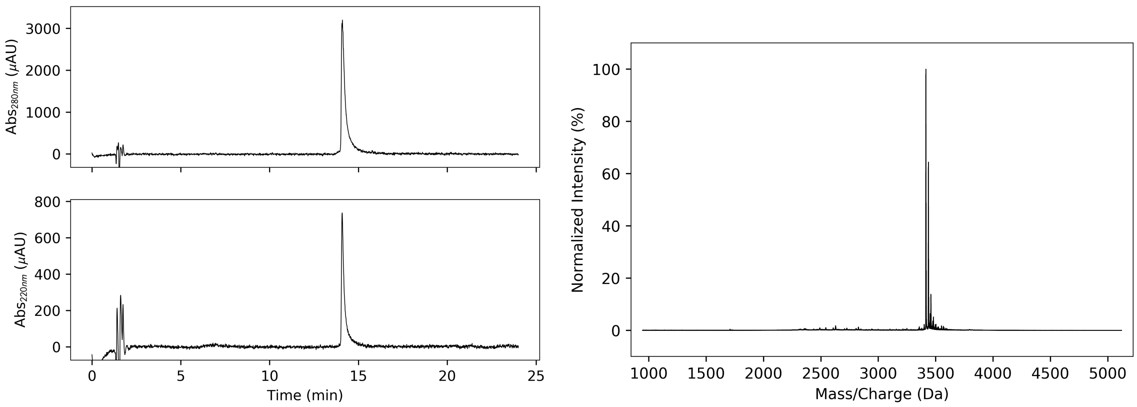


**Figure S 2.14 CC-Type2-$(\mathbf{I_aY_d})_4$-Y3F-Y24F** - HPLC traces (left, 220 and 280 nm) and MALDI-TOF MS (right). Calculated mass = 3413.8 Da, observed mass = 3416 Da.

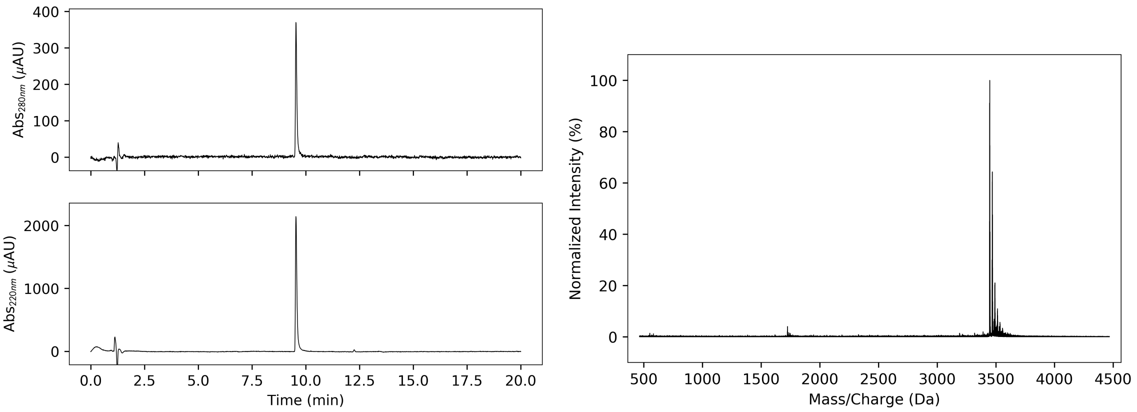


**Figure S 2.15 CC-Type2-($I_aY_d)_4$_ r** - HPLC traces (left, 220 and 280 nm) and MALDI-TOF MS (right). Calculated mass = 3444.8 Da, observed mass = 3446 Da.

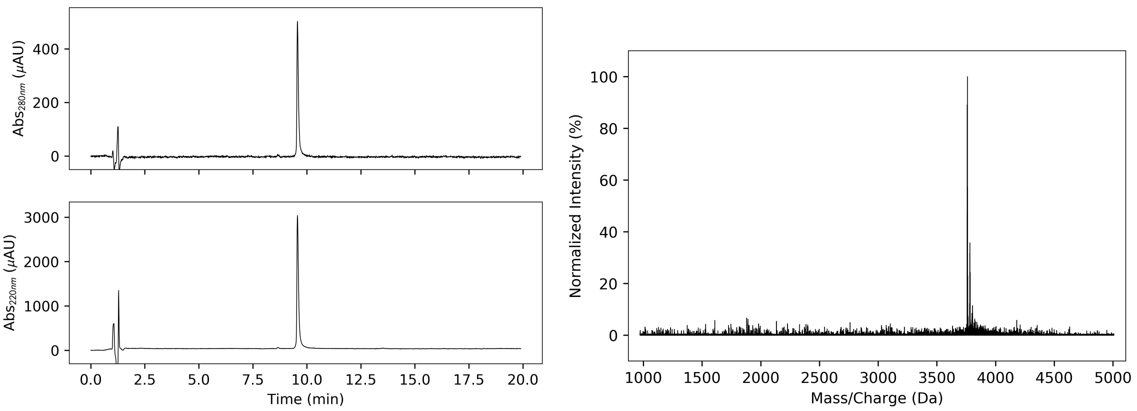


**Figure S 2.16 CC-Type2-($I_aY_d)_{4.5}$** - HPLC traces (left, 220 and 280 nm) and MALDI-TOF MS (right). Calculated mass = 3757.0 Da, observed mass = 3758 Da.

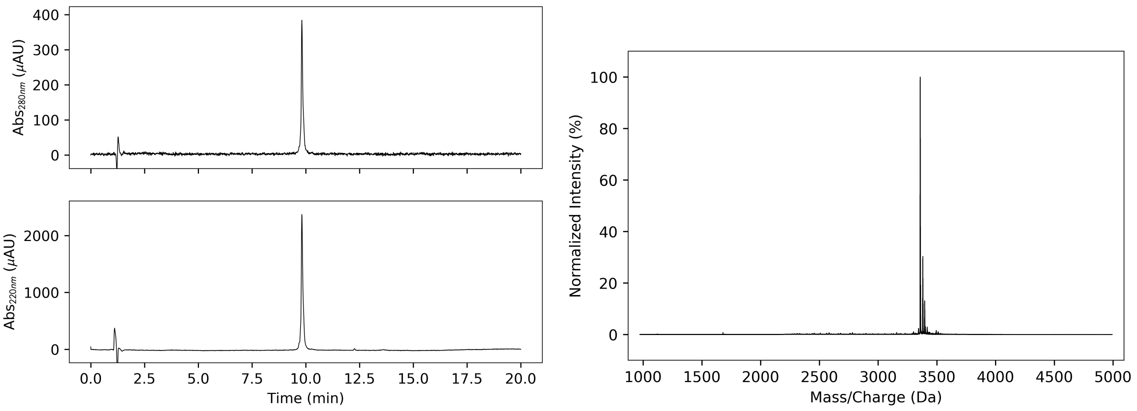


**Figure S 2.17 CC-Type2-($V_aY_d)_4$-Y3F-Y24F** - HPLC traces (left, 220 and 280 nm) and MALDI-TOF MS (right). Calculated mass = 3356.7 Da, observed mass = 3357 Da.

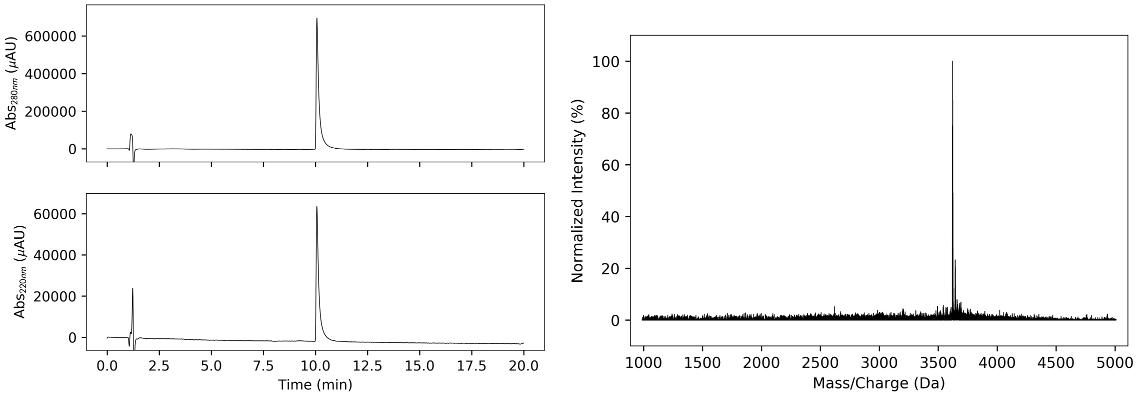


**Figure S 2.18 CC-Type2-($Y_aF_d$)$_4$-W19(BrPhe)** - HPLC traces (left, 220 and 280 nm) and MALDI-TOF MS (right). Calculated mass = 3619.6 Da, observed mass = 3621 Da.

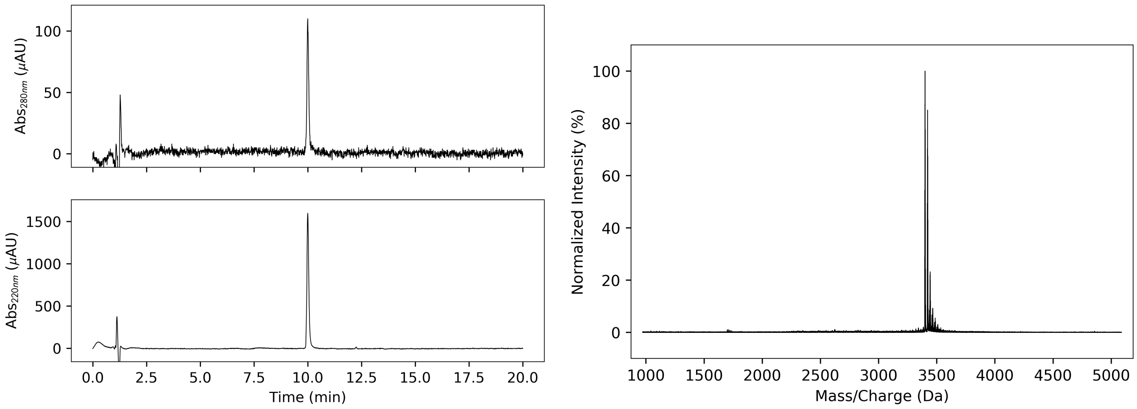


**Figure S 2.19 CC-Type2-($V_aY_d$)$_4$-Y3F-W19(BrPhe)-Y24F** - HPLC traces (left, 220 and 280 nm) and MALDI-TOF MS (right). Calculated mass = 3395.7 Da, observed mass = 3397 Da.

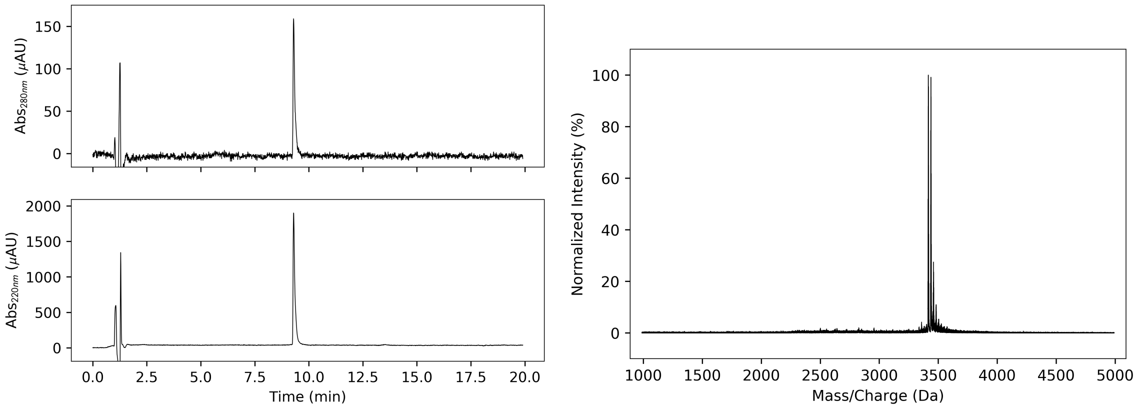


**Figure S 2.20 CC-Type2-($V_aY_d$)$_4$-Y3F-W19(BrPhe)** - HPLC traces (left, 220 and 280 nm) and MALDI-TOF MS (right). Calculated mass = 3411.6 Da, observed mass = 3413 Da.

### 3 Circular dichroism (CD)

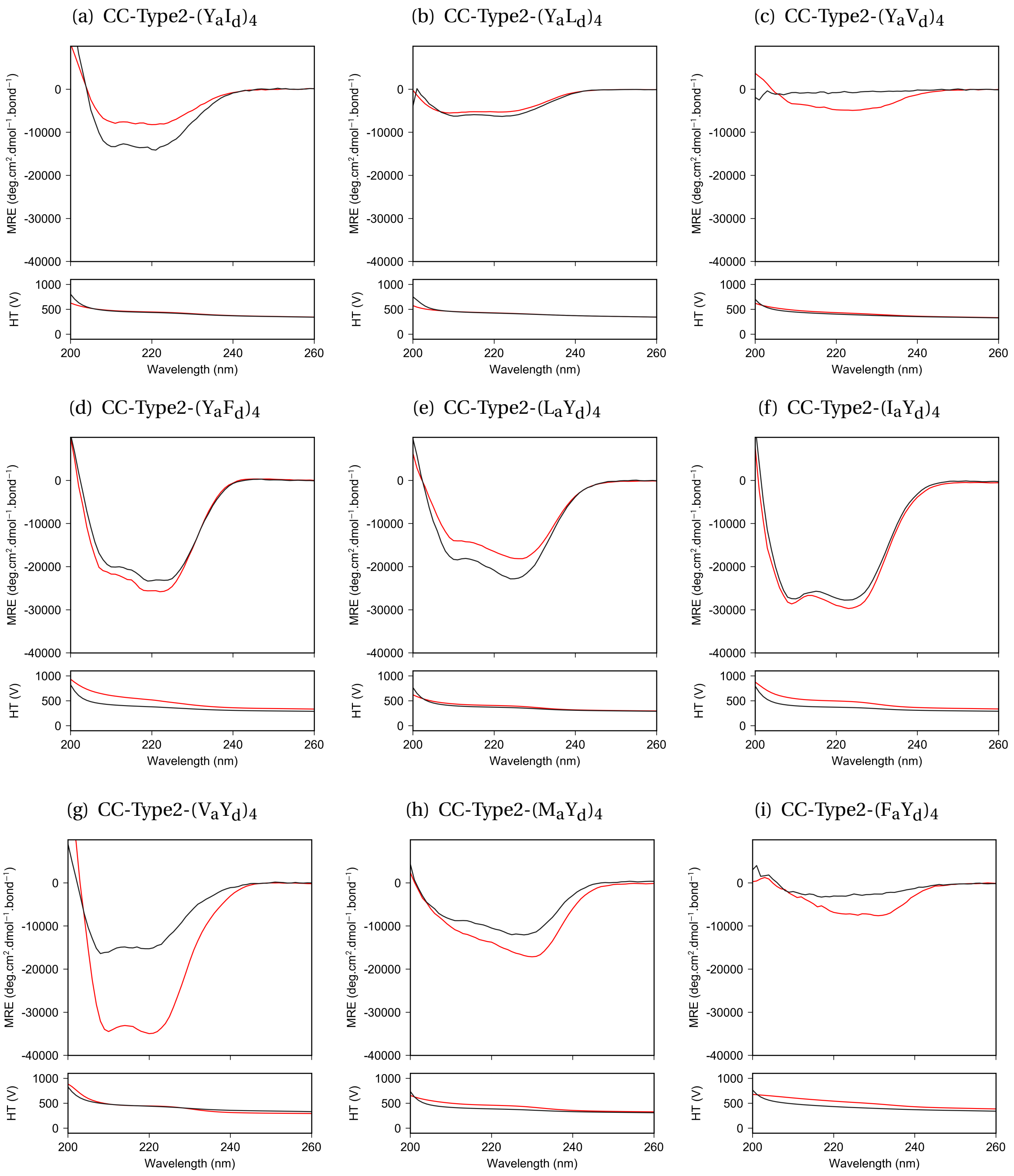


**Figure S 3.1** Screening CD spectra at 20 °C measured at 10 $\mu$M peptide concentration (black) and 100 $\mu$M peptide concentration (red). Conditions: PBS (pH 7.4).

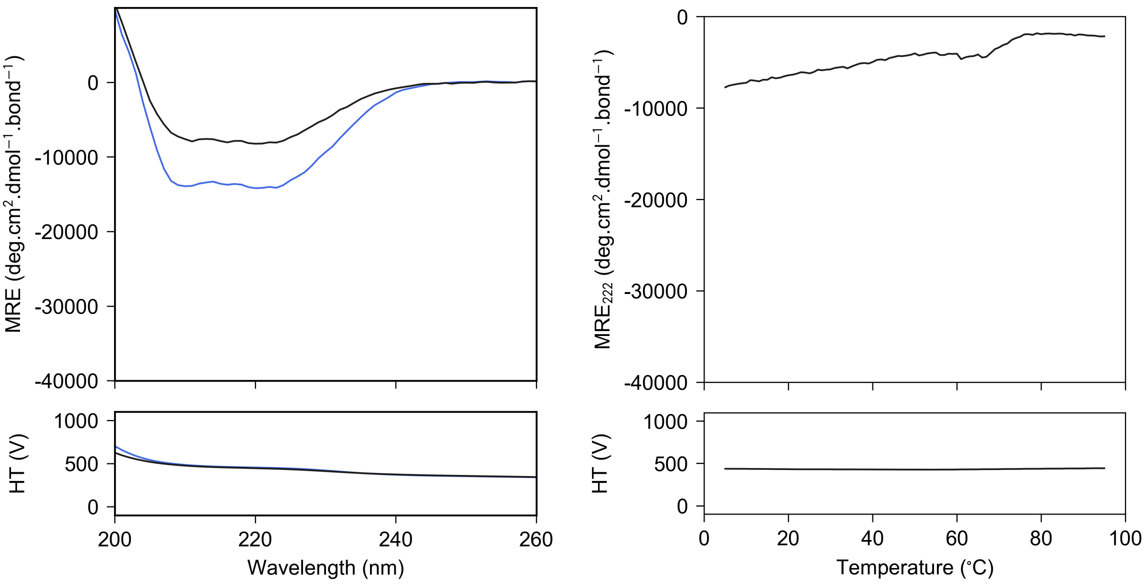


**Figure S 3.2 CC-Type2-($Y_aI_d$)$_4$** - CD spectrum at 20 °C (left) and thermal denaturation profile monitored at 222 nm (right). Conditions: 100 $\mu$M peptide concentration, PBS (pH 7.4).

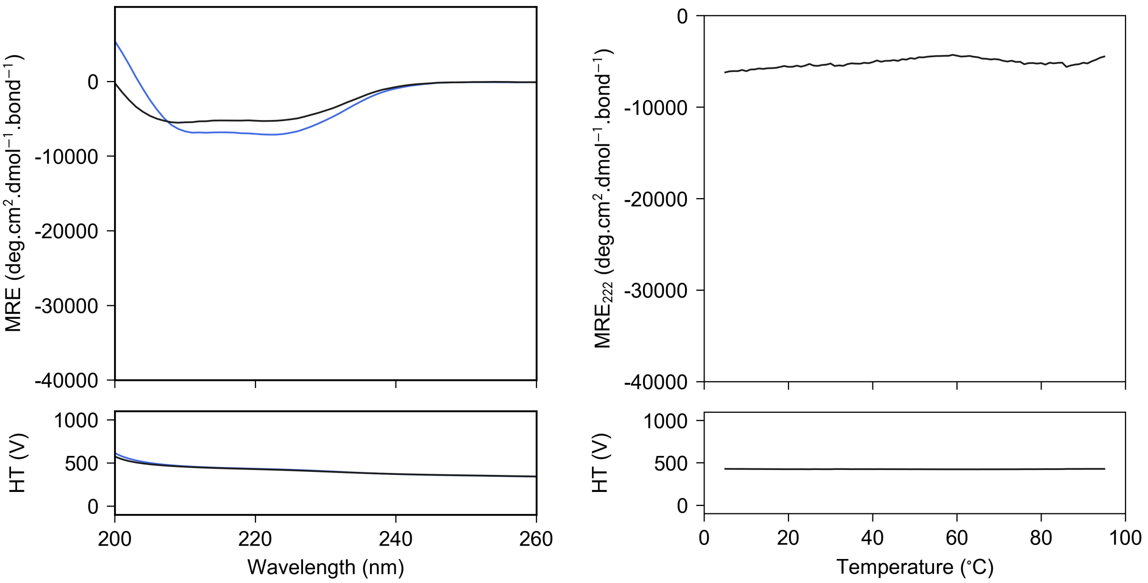


**Figure S 3.3 CC-Type2-($Y_aL_d$)$_4$** - CD spectrum at 20 °C (left) and thermal denaturation profile monitored at 222 nm (right). Conditions: 100 $\mu$M peptide concentration, PBS (pH 7.4).

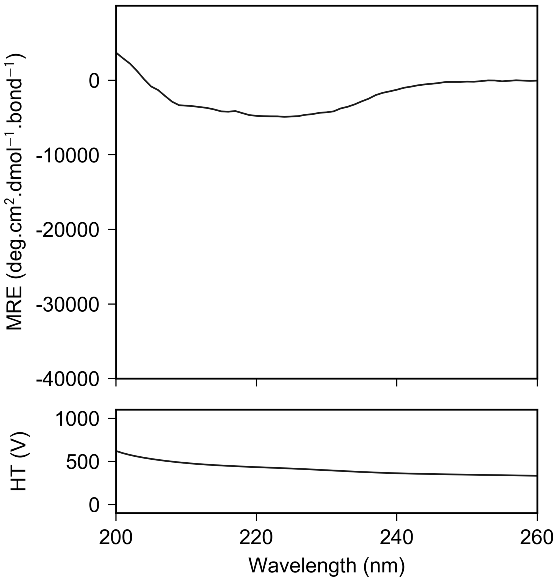


**Figure S 3.4 CC-Type2-($Y_aV_d$)$_4$** - CD spectrum at 20 °C (left). Conditions: 100 $\mu$M peptide concentration, PBS (pH 7.4).

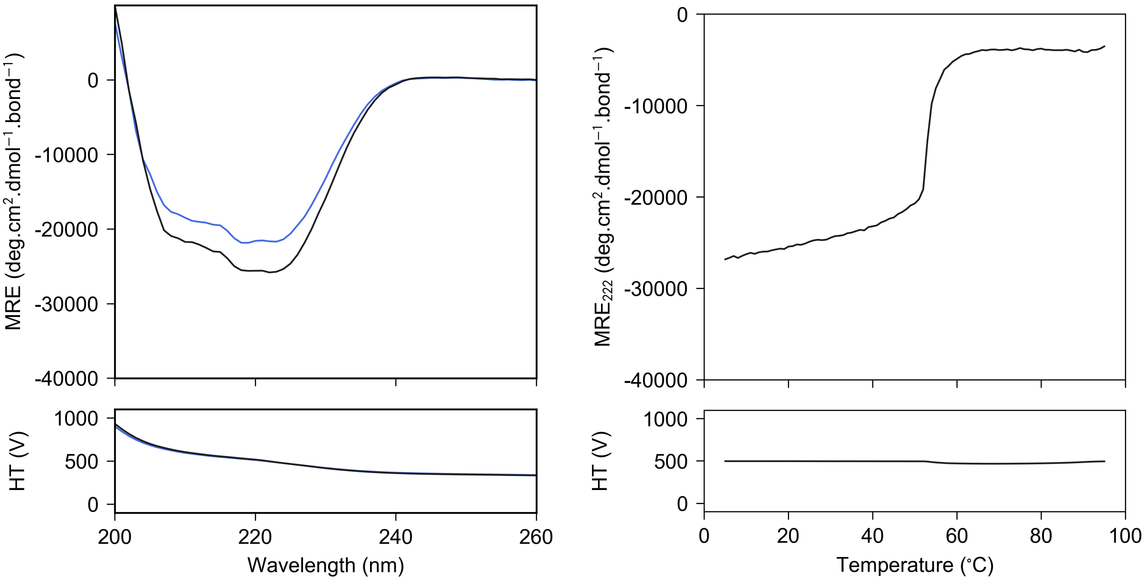


**Figure S 3.5 CC-Type2-($Y_aF_d$)$_4$** - CD spectrum at 20 °C (left) and thermal denaturation profile monitored at 222 nm (right). Conditions: 100 $\mu$M peptide concentration, PBS (pH 7.4).

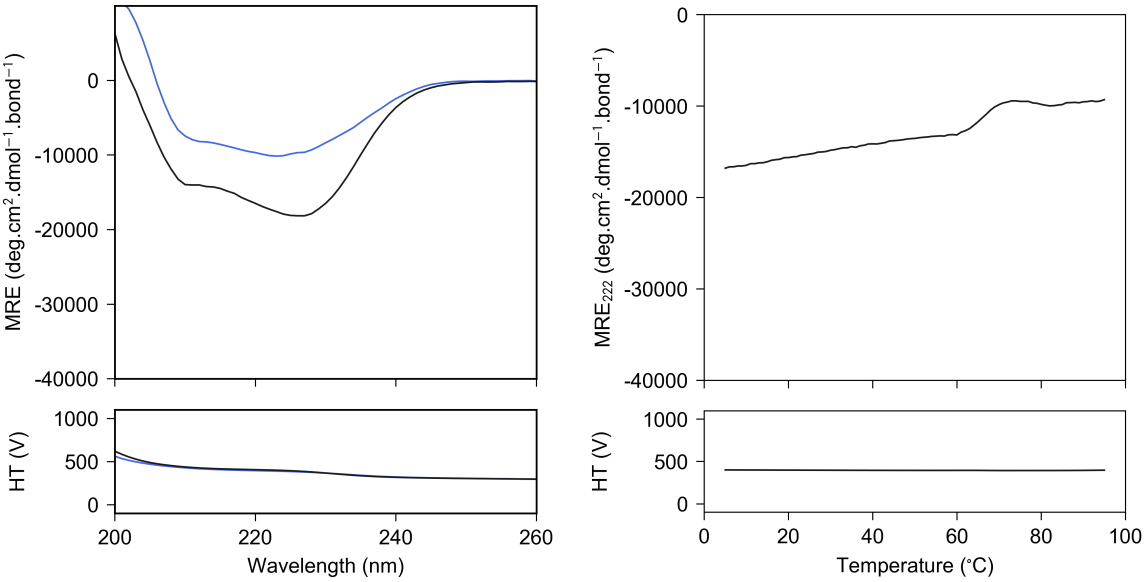


**Figure S 3.6 CC-Type2-($L_aY_d$)$_4$** - CD spectrum at 20 °C (left) and thermal denaturation profile monitored at 222 nm (right). Conditions: 100 $\mu$M peptide concentration, PBS (pH 7.4).

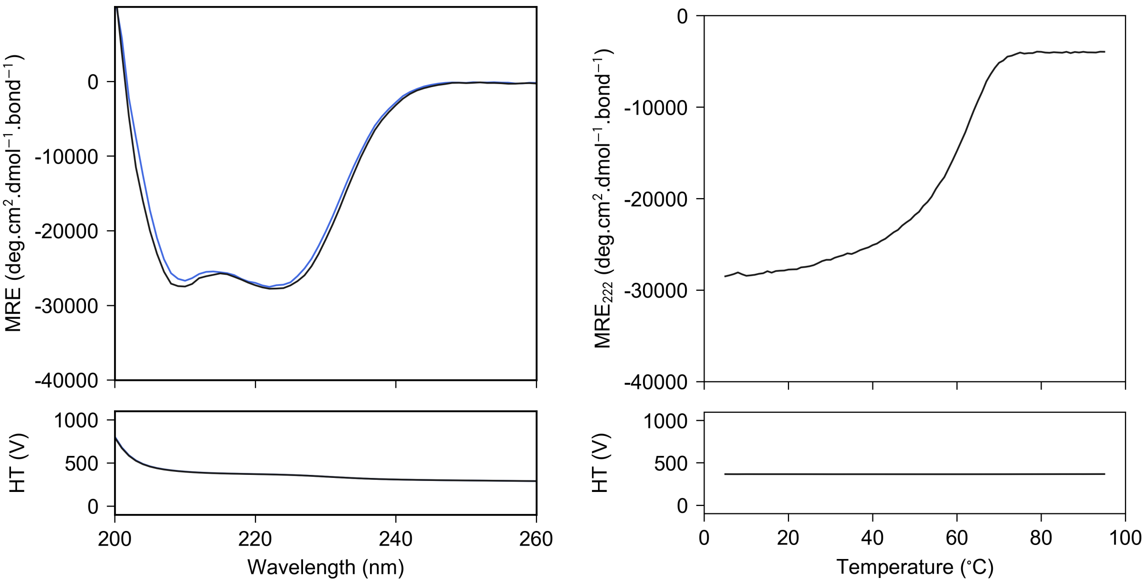


**Figure S 3.7 CC-Type2-($I_aY_d$)$_4$** - CD spectrum at 20 °C (left) and thermal denaturation profile monitored at 222 nm (right). Conditions: 10 $\mu$M peptide concentration, PBS (pH 7.4).

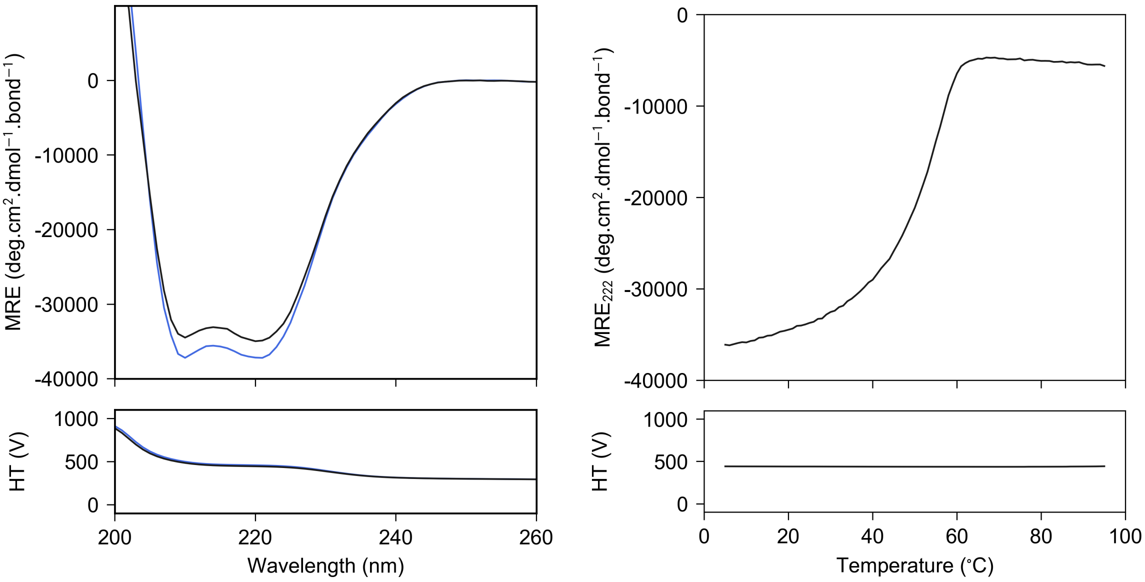


**Figure S 3.8 CC-Type2-($V_aY_d$)$_4$** - CD spectrum at 20 °C (left) and thermal denaturation profile monitored at 222 nm (right). Conditions: 100 $\mu$M peptide concentration, PBS (pH 7.4).

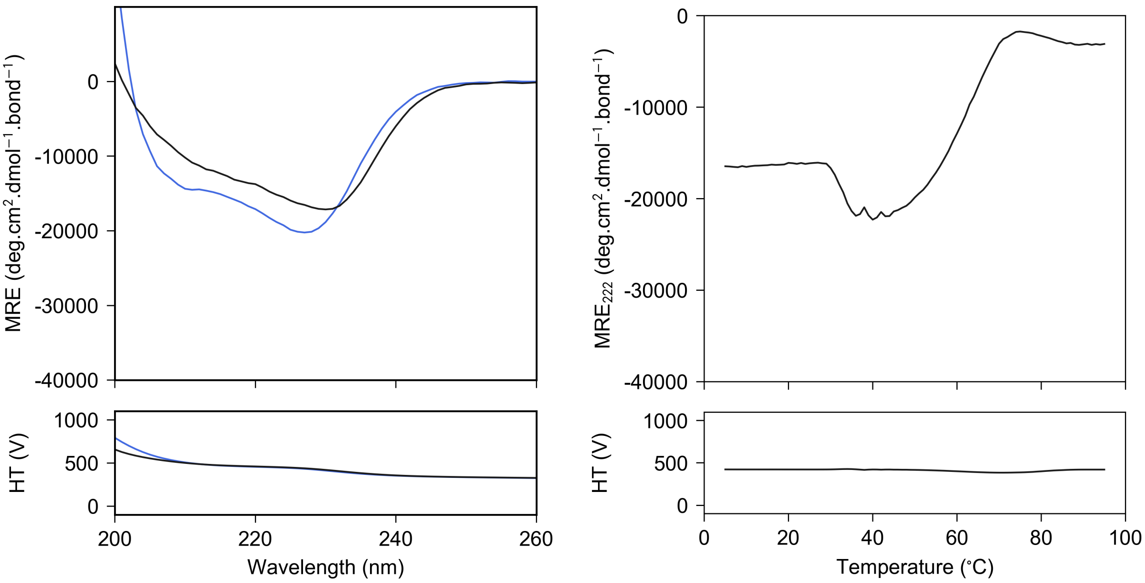


**Figure S 3.9 CC-Type2-($M_aY_d$)$_4$** - CD spectrum at 20 °C (left) and thermal denaturation profile monitored at 222 nm (right). Conditions: 100 $\mu$M peptide concentration, PBS (pH 7.4).

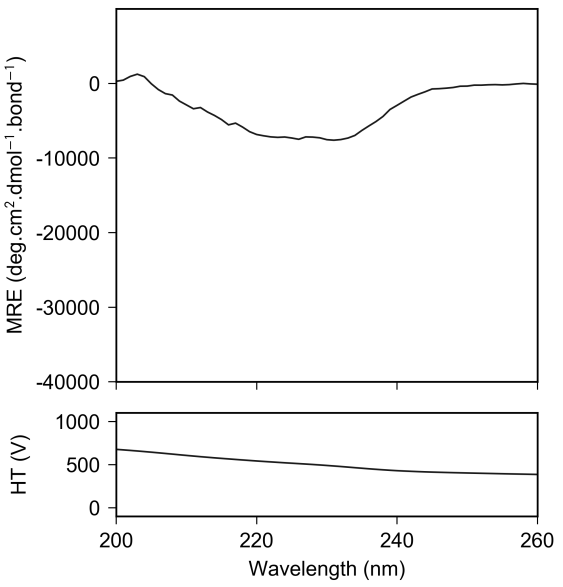


**Figure S 3.10 CC-Type2-($F_aY_d$)$_4$** - CD spectrum at 20 °C (left). Conditions: 100 $\mu$M peptide concentration, PBS (pH 7.4).

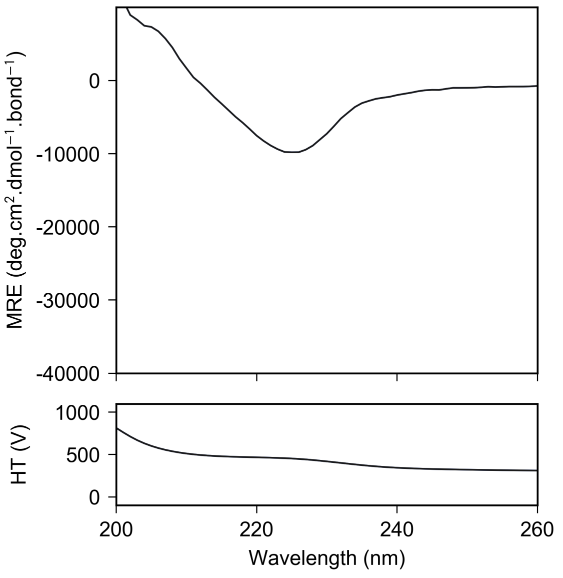


**Figure S 3.11 CC-Type2-($Y_aY_d$)$_4$** - CD spectrum at 20 °C (left). Conditions: 100 $\mu$M peptide concentration, PBS (pH 7.4).

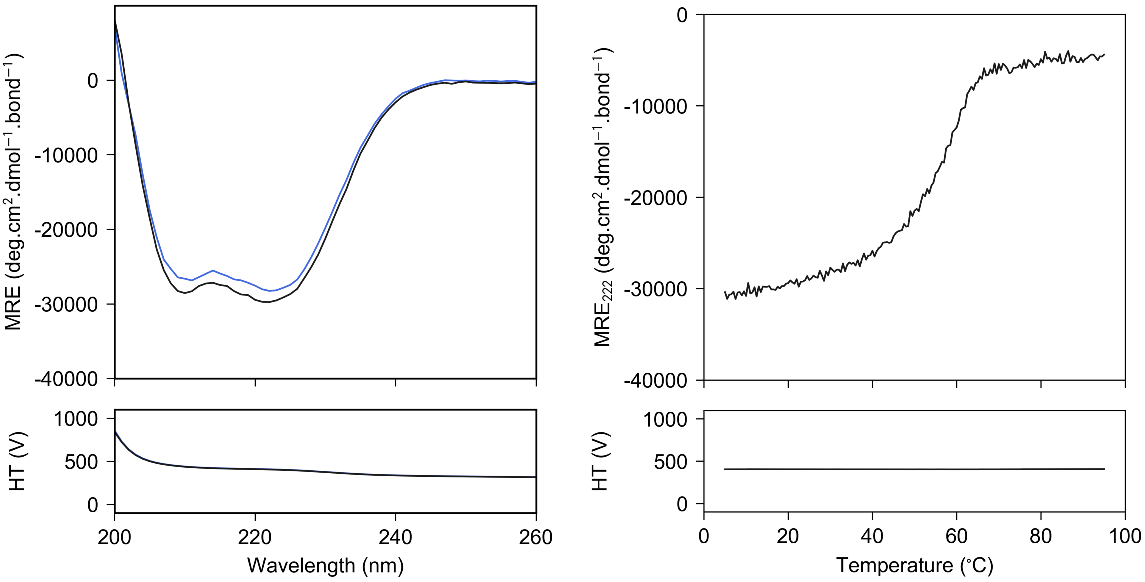


**Figure S 3.12 CC-Type2-($I_aY_d$)$_4$_ termK** - CD spectrum at 20 °C (left) and thermal denaturation profile monitored at 222 nm (right). Conditions: 10 $\mu$M peptide concentration, PBS (pH 7.4).

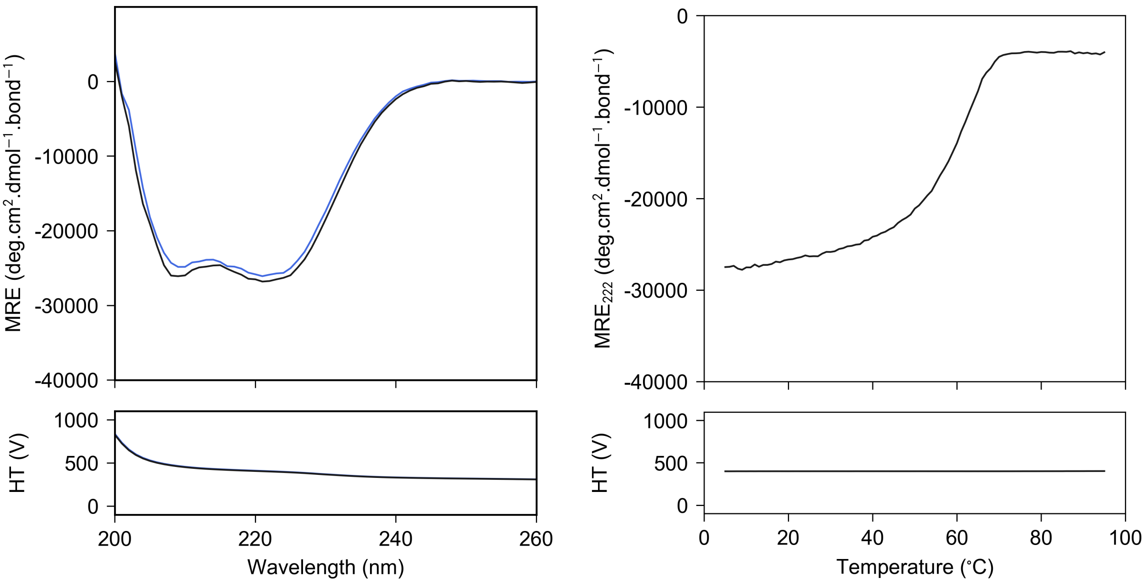


**Figure S 3.13 CC-Type2-($I_aY_d$)$_4$_ termSS** - CD spectrum at 20 °C (left) and thermal denaturation profile monitored at 222 nm (right). Conditions: 10 $\mu$M peptide concentration, PBS (pH 7.4).

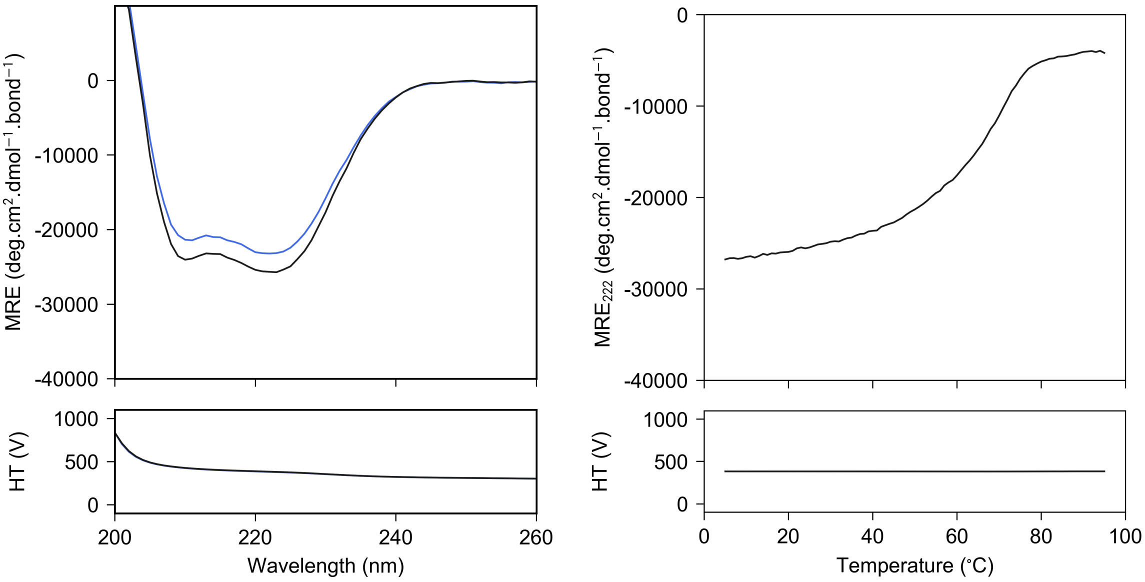


**Figure S 3.14 CC-Type2-($I_aY_d$)$_4$-Y3F** - CD spectrum at 20 °C (left) and thermal denaturation profile monitored at 222 nm (right). Conditions: 10 $\mu$M peptide concentration, PBS (pH 7.4).

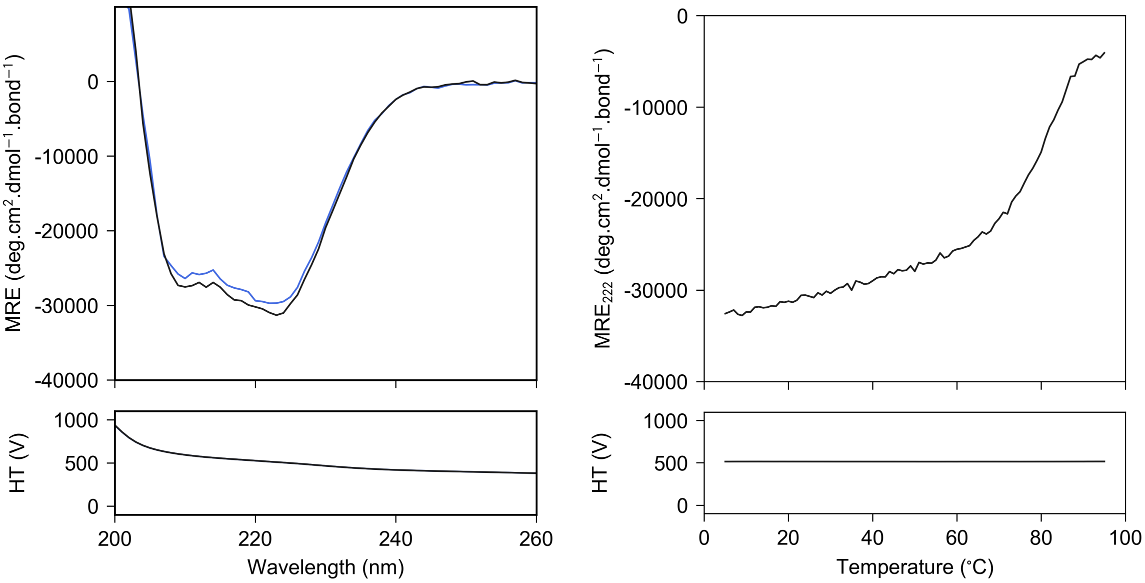


**Figure S 3.15 CC-Type2-($I_aY_d$)$_4$-Y3F-Y24F** - CD spectrum at 20 °C (left) and thermal denaturation profile monitored at 222 nm (right). Conditions: 10 $\mu$M peptide concentration, PBS (pH 7.4).

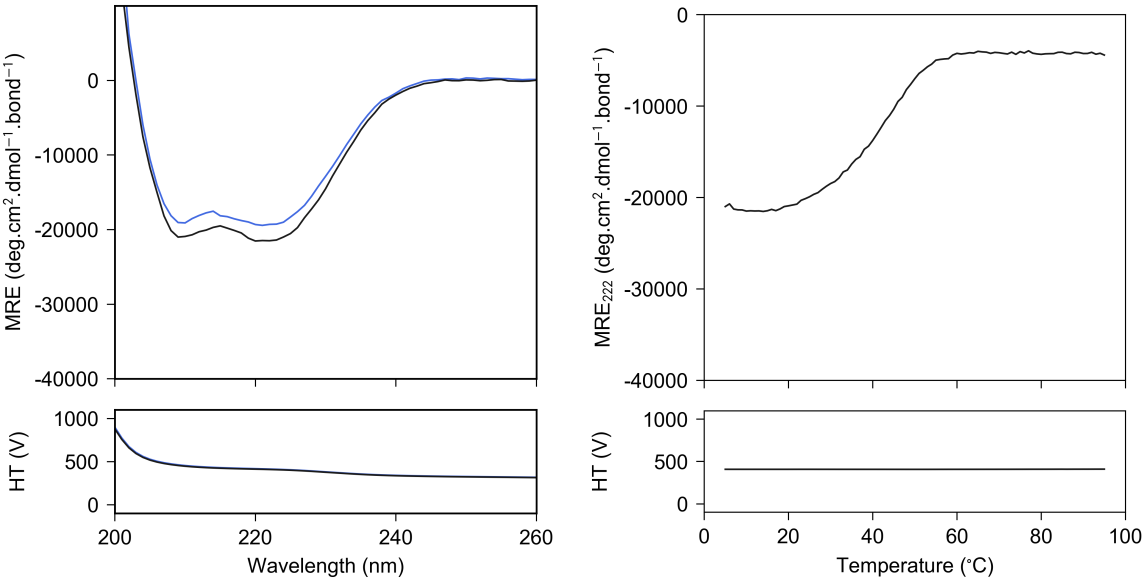


**Figure S 3.16 CC-Type2-$(I_aY_d)_{4\_}$ r** - CD spectrum at 20 °C (left) and thermal denaturation profile monitored at 222 nm (right). Conditions: 10 $\mu$M peptide concentration, PBS (pH 7.4).

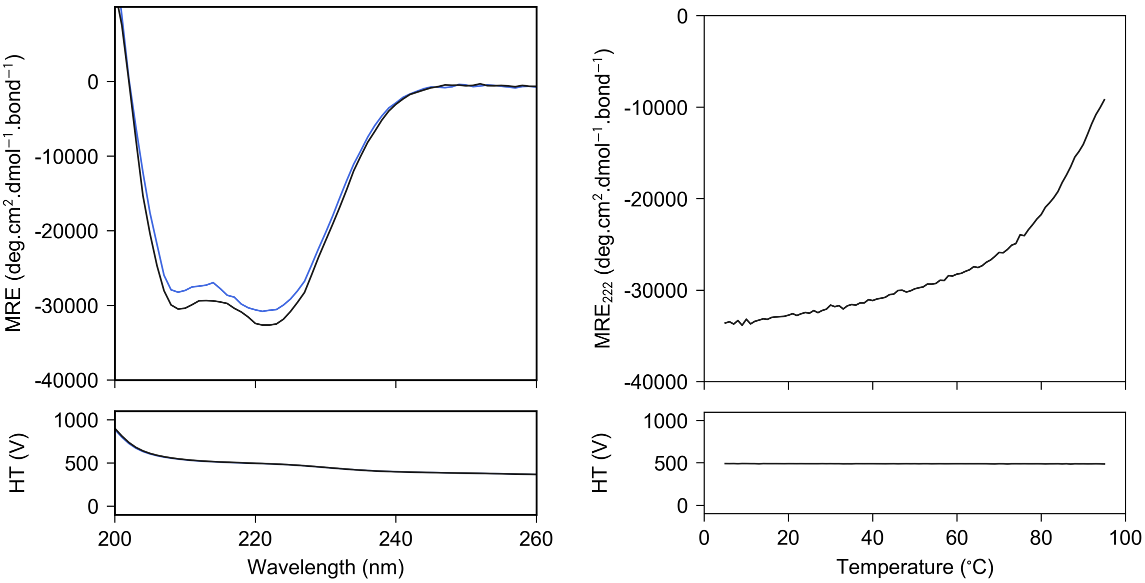


**Figure S 3.17 CC-Type2-$(I_aY_d)_{4.5}$** - CD spectrum at 20 °C (left) and thermal denaturation profile monitored at 222 nm (right). Conditions: 10 $\mu$M peptide concentration, PBS (pH 7.4).

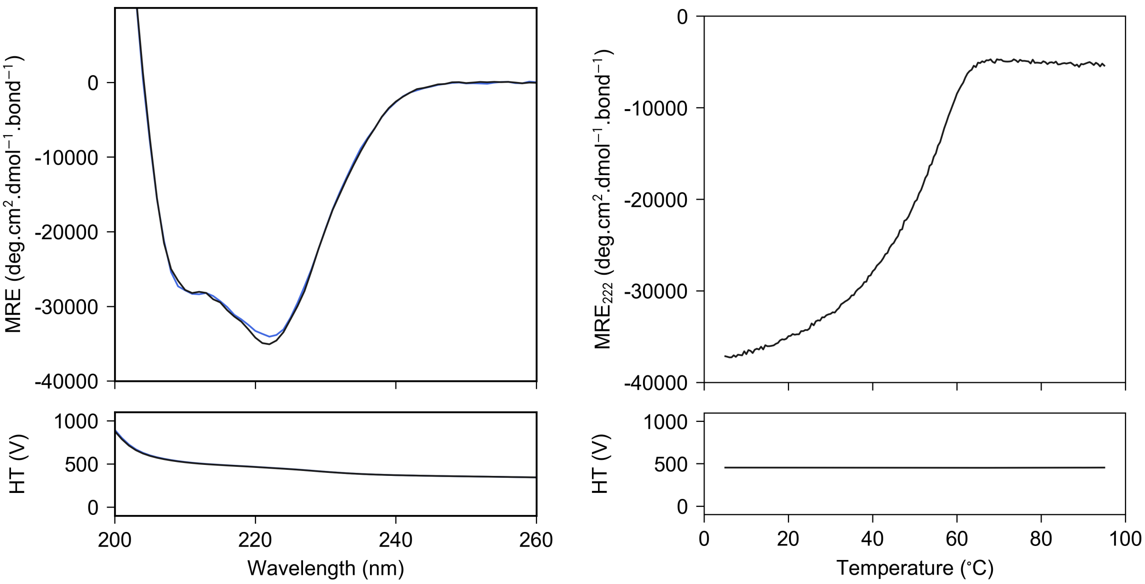


**Figure S 3.18 CC-Type2-($V_aY_d$)$_4$-Y3F-Y24F** - CD spectrum at 20 °C (left) and thermal denaturation profile monitored at 222 nm (right). Conditions: 10 $\mu$M peptide concentration, PBS (pH 7.4).

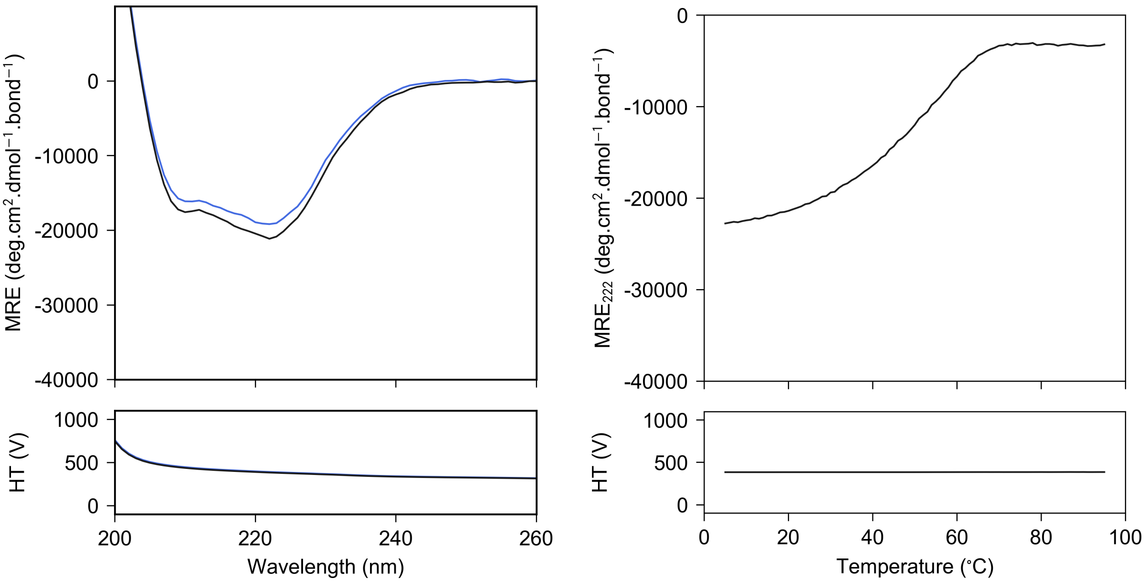


**Figure S 3.19 CC-Type2-($V_aY_d$)$_4$-Y3F-W19(BrPhe)-Y24F** - CD spectrum at 20 °C (left) and thermal denaturation profile monitored at 222 nm (right). Conditions: 10 $\mu$M peptide concentration, PBS (pH 7.4).

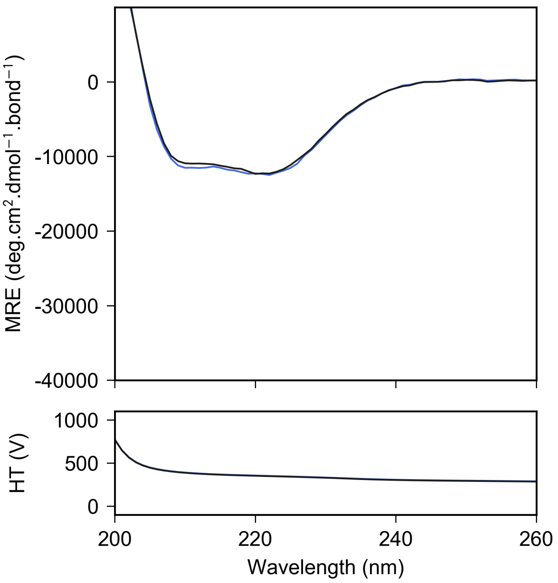


**Figure S 3.20 CC-Type2-($V_aY_d$)$_4$-Y3F-W19(BrPhe)** - CD spectrum at 20 °C (left) and thermal denaturation profile monitored at 222 nm (right). Conditions: 10 $\mu$M peptide concentration, PBS (pH 7.4).

### 4 Analytical ultracentrifuge (AUC)

Due to low $\alpha$ helicity, the AUC experiments were not conducted for the sequences CC-Type2-$(Y_aI_d)_4$ and CC-Type2-$(Y_aL_d)_4$. CC-Type2-$(Y_aV_d)_4$, CC-Type2-$(L_aY_d)_4$, CC-Type2-$(M_aY_d)_4$ and CC-Type2-$(F_aY_d)_4$ aggregated at 3,000 rpm. Large distributions of species were observed during c(s) distribution analysis for CC-Type2-$(I_aY_d)_4$, CC-Type2-$(V_aY_d)_4$, CC-Type2-$(F_aY_d)_4$, CC-Type2-$(I_aY_d)_4$_ termK and CC-Type2-$(I_aY_d)_4$_ termSS. The c(s) distribution of CC-Type2-$(I_aY_d)_4$ is provided as an example.

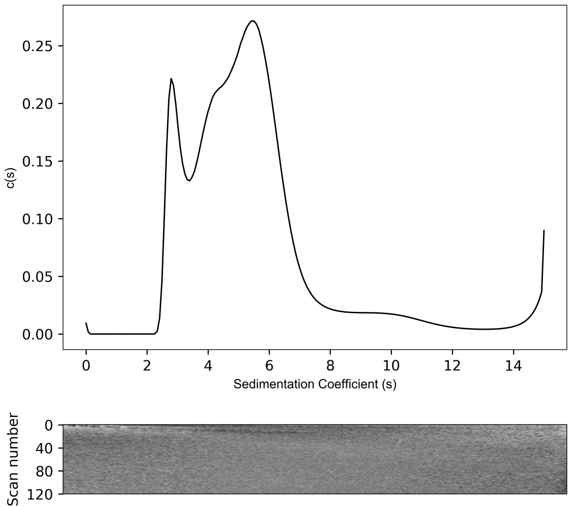


**Figure S 4.1 CC-Type2-$(I_aY_d)_4$** - AUC data and fits (top) and residuals (bottom) ($\bar{\upsilon}$ = 0.748 $cm^3$ $g^{-1}$). Continuous c(s) distribution from sedimentation-velocity data at 50k rpm returning a large distribution of masses. Conditions: 150 $\mu$M peptide concentration, PBS (pH 7.4).

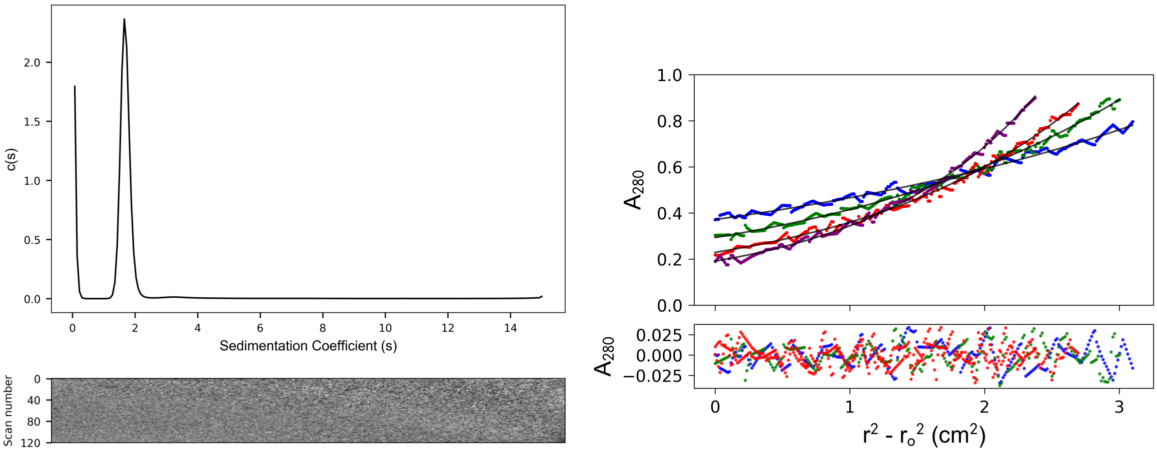


**Figure S 4.2 CC-Type2-($Y_aF_d$)$_4$** - AUC data and fits (top) and residuals (bottom) ($\bar{v}$ = 0.732 cm$^3$ g$^{-1}$). Left: continuous c(s) distribution from sedimentation-velocity data at 50k rpm returning s = 1.689 S, s20,w = 1.751 S, f/f0 = 1.250 and mw = 15,219 Da (4.3 × monomer mass) at 95% confidence level. Conditions: 150 $\mu$M peptide concentration, PBS (pH 7.4). Right: sedimentation-equilibrium data (top, dots) and fitted single-ideal species model curves at 18k (blue), 22k (red), 26k (green) and 30k (purple) rpm. The fit returns a mass of 15,530 Da (4.3 × monomer mass, 95% confidence limits 15,466 – 15,629). Bottom: residuals for the above fits using the same colour scheme as above. Conditions: 150 $\mu$M peptide concentration, PBS (pH 7.4).

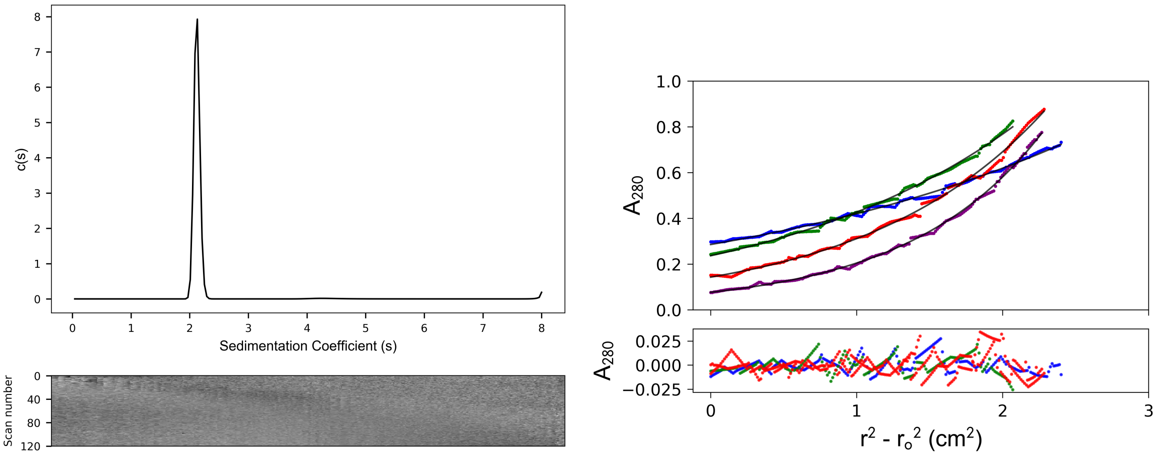


**Figure S 4.3 CC-Type2-($I_aY_d$)$_4$-Y3F** - AUC data and fits (top) and residuals (bottom) ($\bar{v}$ = 0.759 cm$^3$ g$^{-1}$). Left: continuous c(s) distribution from sedimentation-velocity data at 50k rpm returning s = 2.123 S, s20,w = 2.347 S, f/f0 = 1.177 and mw = 21,510 Da (6.3 × monomer mass) at 95% confidence level. Conditions: 150 $\mu$M peptide concentration, PBS (pH 7.4). Right: sedimentation-equilibrium data (top, dots) and fitted single-ideal species model curves at 18k (blue), 22k (red), 26k (green) and 30k (purple) rpm. The fit returns a mass of 21,750 Da (6.3 × monomer mass, 95% confidence limits 21,601 – 21,960). Bottom: residuals for the above fits using the same colour scheme as above. Conditions: 150 $\mu$M peptide concentration, PBS (pH 7.4).

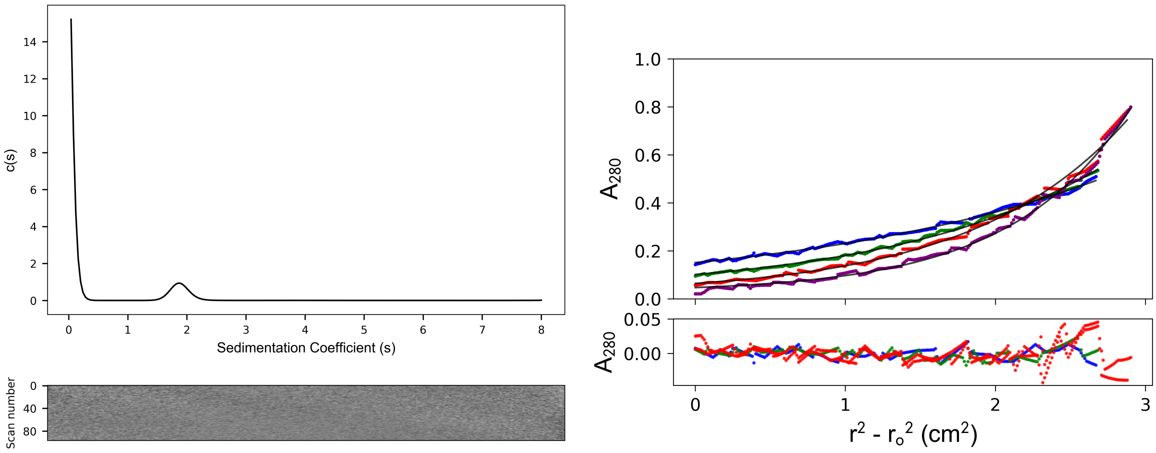


**Figure S 4.4 CC-Type2-$(I_aY_d)_4$-Y3F-Y24F** - AUC data and fits (top) and residuals (bottom) ($\bar{\upsilon}$ = 0.753 $cm^3$ $g^{-1}$). Left: continuous c(s) distribution from sedimentation-velocity data at 50k rpm returning s = 1.882 S, s20,w = 2.141 S, f/f0 = 1.253 and mw = 20,867 Da (6.1 × monomer mass) at 95% confidence level. Conditions: 150 $\mu$M peptide concentration, PBS (pH 7.4). Right: sedimentation-equilibrium data (top, dots) and fitted single-ideal species model curves at 20k (blue), 24k (red), 28k (green) and 32k (purple) rpm. The fit returns a mass of 22,570 Da (6.6 × monomer mass, 95% confidence limits 22,267 – 22,885). Bottom: residuals for the above fits using the same colour scheme as above. Conditions: 150 $\mu$M peptide concentration, PBS (pH 7.4).

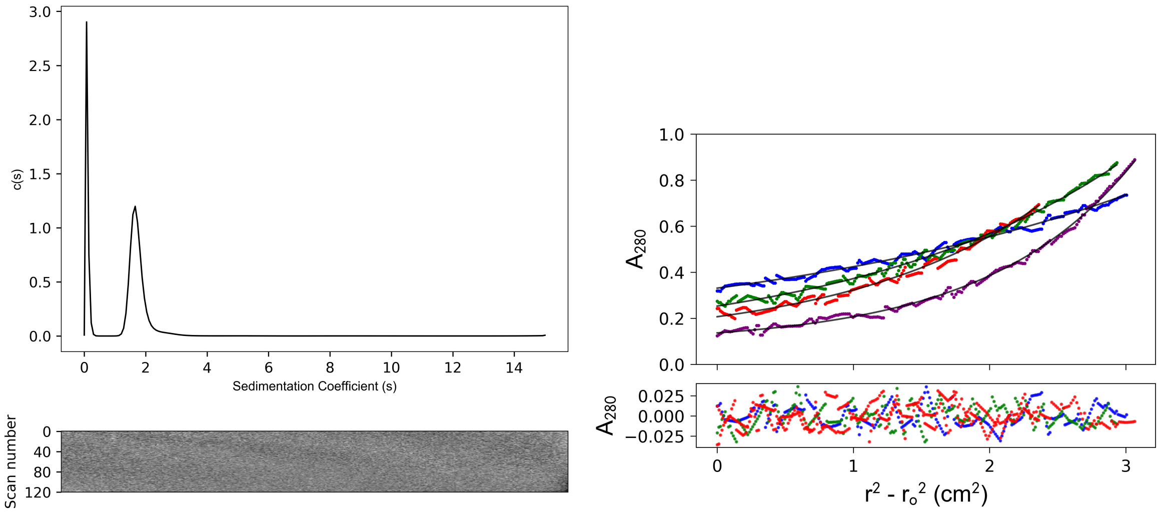


**Figure S 4.5 CC-Type2-$(I_aY_d)_4$_ r** - AUC data and fits (top) and residuals (bottom) ($\bar{\upsilon}$ = 0.748 $cm^3$ $g^{-1}$). Left: continuous c(s) distribution from sedimentation-velocity data at 50k rpm returning s = 1.733 S, s20,w = 1.800 S, f/f0 = 1.418 and mw = 21,086 Da (6.1 × monomer mass) at 95% confidence level. Conditions: 150 $\mu$M peptide concentration, PBS (pH 7.4). Right: sedimentation-equilibrium data (top, dots) and fitted single-ideal species model curves at 18k (blue), 22k (red), 26k (green) and 30k (purple) rpm. The fit returns a mass of 20,430 Da (5.9 × monomer mass, 95% confidence limits 20,191 – 20,523). Bottom: residuals for the above fits using the same colour scheme as above. Conditions: 150 $\mu$M peptide concentration, PBS (pH 7.4).

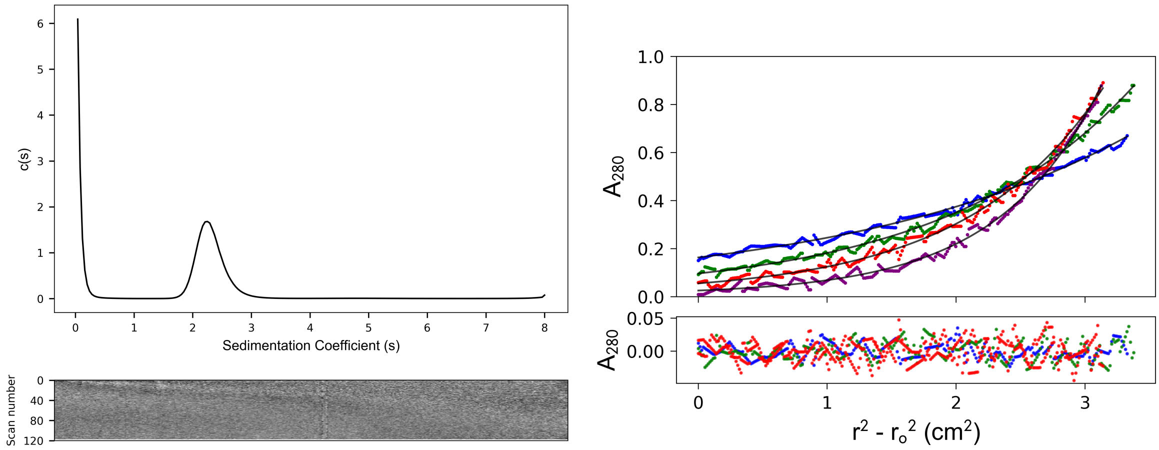


**Figure S 4.6 CC-Type2-($I_aY_d$)$_{4.5}$** - AUC data and fits (top) and residuals (bottom) ($\bar{\upsilon}$ = 0.755 $cm^3$ $g^{-1}$). Left: continuous c(s) distribution from sedimentation-velocity data at 50k rpm returning s = 2.304 S, s20,w = 2.616 S, f/f0 = 1.102 and mw = 22,984 Da (6.1 × monomer mass) at 95% confidence level. Conditions: 150 $\mu$M peptide concentration, PBS (pH 7.4). Right: sedimentation-equilibrium data (top, dots) and fitted single-ideal species model curves at 18k (blue), 22k (red), 26k (green) and 30k (purple) rpm. The fit returns a mass of 27,300 Da (7.3 × monomer mass, 95% confidence limits 26,466 – 28,717). Bottom: residuals for the above fits using the same colour scheme as above. Conditions: 150 $\mu$M peptide concentration, PBS (pH 7.4).

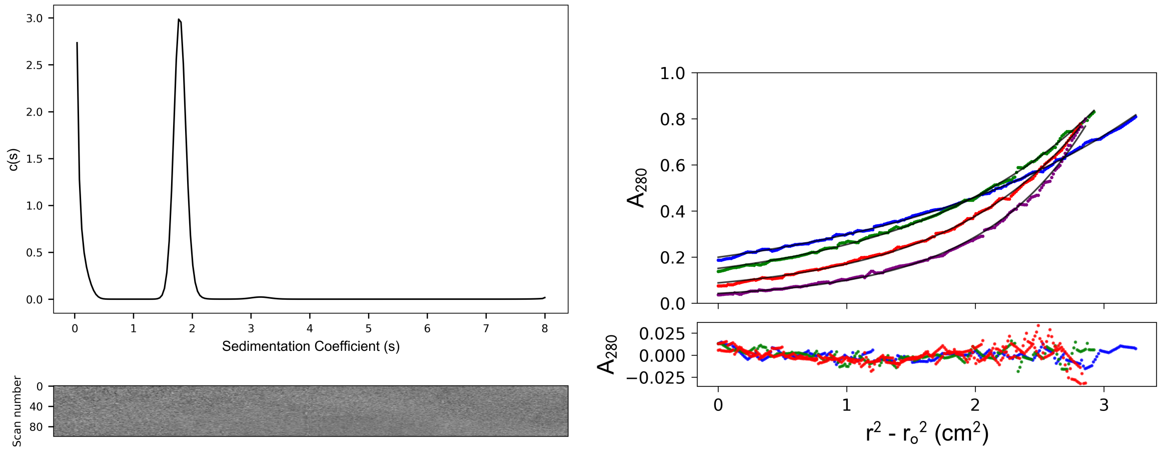


**Figure S 4.7 CC-Type2-($V_aY_d$)$_4$-Y3F-Y24F** - AUC data and fits (top) and residuals (bottom) ($\bar{\upsilon}$ = 0.746 $cm^3$ $g^{-1}$). Left: continuous c(s) distribution from sedimentation-velocity data at 50k rpm returning s = 1.793 S, s20,w = 1.980 S, f/f0 = 1.244 and mw = 18,036 Da (5.4 × monomer mass) at 95% confidence level. Conditions: 150 $\mu$M peptide concentration, PBS (pH 7.4). Right: sedimentation-equilibrium data (top, dots) and fitted single-ideal species model curves at 22k (blue), 26k (red), 30k (green) and 34k (purple) rpm. The fit returns a mass of 18,320 Da (5.5 × monomer mass, 95% confidence limits 18,162 – 18,484). Bottom: residuals for the above fits using the same colour scheme as above. Conditions: 150 $\mu$M peptide concentration, PBS (pH 7.4).

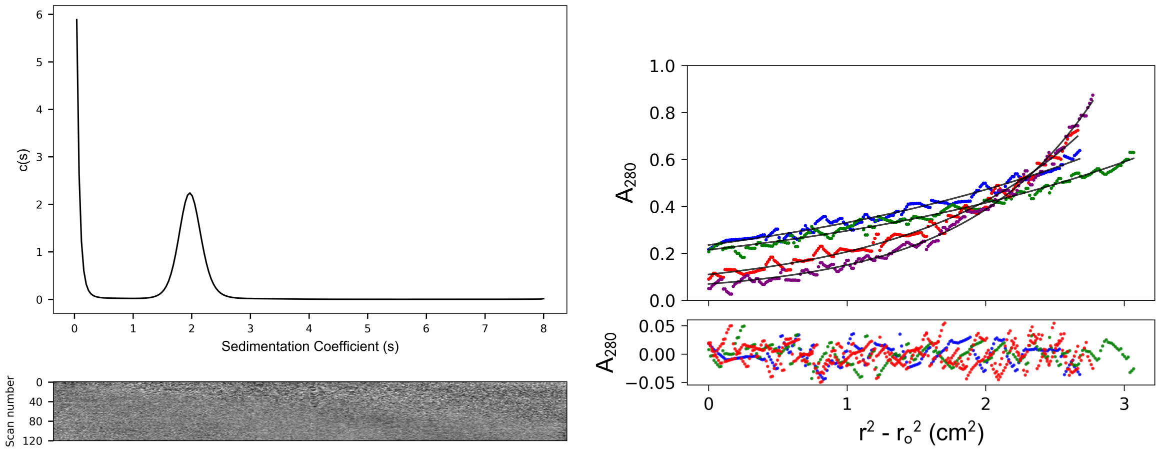


**Figure S 4.8 CC-Type2-($V_aY_d$)$_4$-Y3F-W19(BrPhe)-Y24F** - AUC data and fits (top) and residuals (bottom) ($\bar{\upsilon}$ = 0.745 cm$^3$ g$^{-1}$). Left: continuous c(s) distribution from sedimentation-velocity data at 50k rpm returning s = 1.977 S, s20,w = 2.156 S, f/f0 = 1.292 and mw = 21,682 Da (6.4 × monomer mass) at 95% confidence level. Conditions: 150 $\mu$M peptide concentration, PBS (pH 7.4). Right: sedimentation-equilibrium data (top, dots) and fitted single-ideal species model curves at 18k (blue), 22k (red), 26k (green) and 30k (purple) rpm. The fit returns a mass of 19,250 Da (5.7 × monomer mass, 95% confidence limits 19,169 – 19,268). Bottom: residuals for the above fits using the same colour scheme as above. Conditions: 150 $\mu$M peptide concentration, PBS (pH 7.4).

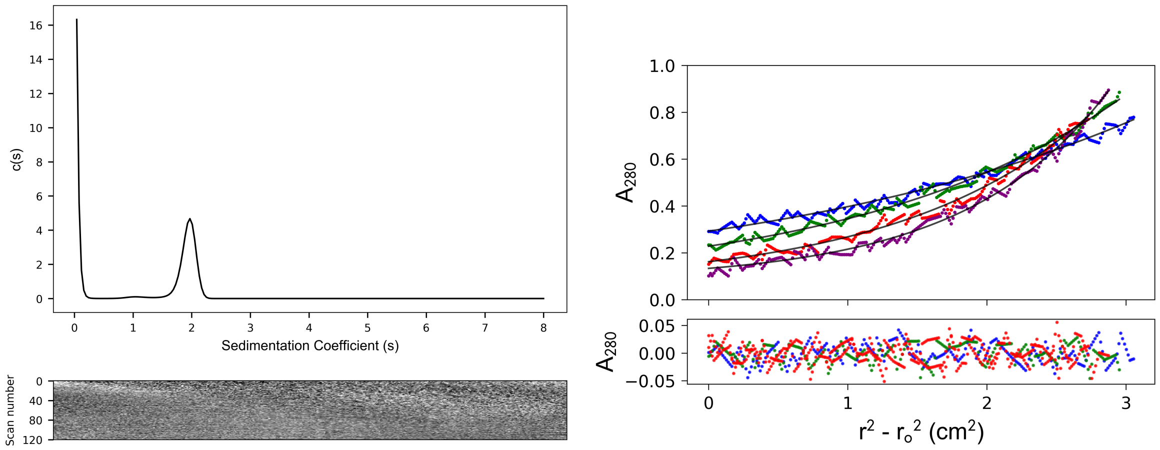


**Figure S 4.9 CC-Type2-($V_aY_d$)$_4$-Y3F-W19(BrPhe)** - AUC data and fits (top) and residuals (bottom) ($\bar{\upsilon}$ = 0.742 cm$^3$ g$^{-1}$). Left: continuous c(s) distribution from sedimentation-velocity data at 50k rpm returning s = 1.939 S, s20,w = 2.090 S, f/f0 = 1.276 and mw = 20,279 Da (6.1 × monomer mass) at 95% confidence level. Conditions: 150 $\mu$M peptide concentration, PBS (pH 7.4). Right: sedimentation-equilibrium data (top, dots) and fitted single-ideal species model curves at 18k (blue), 22k (red), 26k (green) and 30k (purple) rpm. The fit returns a mass of 19,590 Da (5.7 × monomer mass, 95% confidence limits 19,544 – 19,695). Bottom: residuals for the above fits using the same colour scheme as above. Conditions: 150 $\mu$M peptide concentration, PBS (pH 7.4).

### 5 Negative-stain transmission electron microscope (TEM)

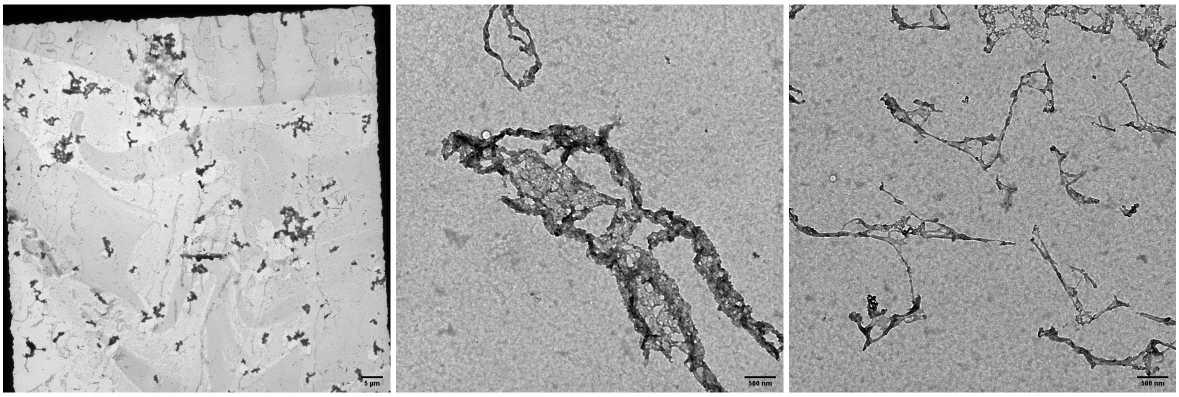


**Figure S 5.1** Transmission electron microscope images of CC-Type2-($I_aY_d$)$_4$, concentrated from a solution at 100 $\mu$M peptide concentration. Conditions: PBS (pH 7.4).

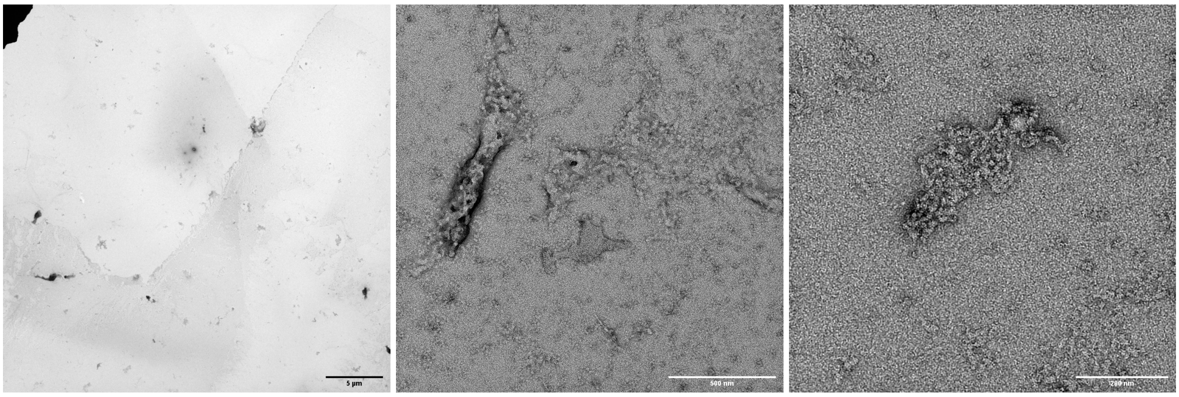


**Figure S 5.2** Transmission electron microscope images of CC-Type2-($V_aY_d$)$_4$, concentrated from a solution at 100 $\mu$M peptide concentration. Conditions: PBS (pH 7.4).

### 6 Crystallography

Crystallisation conditions for each crystal and subsequently, crystallisation data collection and refinement statistics. Crystallisation conditions are final values based on 1:1 dilution with peptide aqueous solution. Highest-resolution shell shown in parentheses and inner shell in square brackets. $R_{free}$ represents the R-factor calculated from 5% of reflections that were not used during refinement.

| Sequence name | Commercial screen | Crystallisation conditions | pH |
|---|---|---|---|
| CC-Type2-$(V_aY_d)_4$-Y3F-W19(BrPhe)-Y24F | Morpheus G10 | 0.05 M Sodium formate, 0.05 M Ammonium acetate, 0.05 M Sodium citrate tribasic dihydrate, 0.05 M Potassium sodium tartrate tetrahydrate, 0.05 M Sodium oxamate, 0.05 M Tris, 0.05 M BICINE, 6% v/v Ethylene glycol and 3 % w/v PEG 8000 | 8.5 |
| CC-Type2-$(V_aY_d)_4$-Y3F-W19(BrPhe) | PACT H8 | 0.1 M sodium sulphate, 0.05 M Bis-Tris propane and 20% w/v PEG 3350 | 8.5 |
| CC-Type2-$(Y_aF_{d)_4}$-W19(BrPhe) | Proplex A8 | 0.1 M Ammonium sulphate, 0.05 M Sodium acetate and 5% w/v PEG 2000 monomethyl ether | 5.5 |

| | CC-Type2-($V_aY_d$)$_4$-Y3F-W19(BrPhe)-Y24F | CC-Type2-($V_aY_d$)$_4$-Y3F-W19(BrPhe) | CC-Type2-($Y_aF_d$)$_4$-W19(BrPhe) |
|---|---|---|---|
| Wavelength (Å) | 0.9198 | 0.9159 | 0.9199 |
| Resolution range (Å) | 38.57–1.84 [38.57–8.23] (1.89–1.84) | 50.90–1.56 [50.90-6.98] (1.60–1.56) | 75.57–1.65 [75.57–7.38] (1.69–1.65) |
| Space group | P 21 21 21 | P 21 21 21 | P 21 21 21 |
| Unit cell dimensions (Å) | 31.15 53.25 111.88 | 44.63 50.9 69.43 | 40.71 75.61 88.12 |
| Unit cell angles (°) | 90 90 90 | 90 90 90 | 90 90 90 |
| Total reflections | 541627 [6642] (15564) | 293806 [3145] (19774) | 439908 [5089] (33190) |
| Unique reflections | 16884 [237] (1199) | 23181 [314] (1672) | 33544 [457] (2442) |
| Multiplicity | 32.1 [28.0] (13.0) | 12.7 [10] (11.8) | 13.1 [11.1] (13.6) |
| Completeness (%) | 99.9 [97.7] (100) | 99.8 [97.9] (99.5) | 100 [99.9] (100) |
| Mean I/sigma(I) | 11.3 [29.2] (0.8) | 12.6 [31.2] (1.3) | 8.7 [21.5] (1.1) |
| Wilson B-factor (Å$^2$) | 45.32 | 25.21 | 22.97 |
| R-merge | 0.176 [0.113] (2.209) | 0.101 [0.065] (1.975) | 0.125 [0.048] (1.968) |
| R-meas | 0.176 [0.115] (2.299) | 0.105 [0.069] (2.064) | 0.13 [0.05] (2.04) |
| R-pim | 0.030 [0.021] (0.632) | 0.030 [0.021] (0.597) | 0.036 [0.015] (0.548) |
| $CC_{1/2}$ | 0.998 [0.997] (0.61) | 0.998 [0.997] (0.609) | 1.0 [1.0] (0.792) |
| $CC^*$ | 1 (0.861) | 1 (0.867) | 0.999 (0.952) |
| Reflections used in refinement | 16834 (1637) | 23130 (2274) | 33364 (3277) |
| Reflections used for R-free | 823 (68) | 1161 (119) | 1702 (196) |
| R-work | 0.2009 (0.3188) | 0.1874 (0.3550) | 0.2338 (0.3842) |
| R-free | 0.2665 (0.2938) | 0.2393 (0.3626) | 0.2641 (0.3949) |
| CC(work) | 0.955 (0.711) | 0.947 (0.741) | 0.932 (0.773) |
| CC(free) | 0.929 (0.683) | 0.936 (0.754) | 0.938 (0.778) |
| Number of non-hydrogen atoms | 1466 | 1598 | 2407 |
| macromolecules | 1407 | 1435 | 2080 |
| ligands | 27 | 17 | 24 |
| solvent | 32 | 146 | 303 |
| Protein residues | 187 | 184 | 250 |
| RMS(bonds) | 0.016 | 0.019 | 0.015 |
| RMS(angles) | 2.02 | 2.06 | 1.70 |
| Ramachandran favored (% ) | 99.33 | 99.33 | 99.50 |
| Ramachandran allowed (% ) | 0.67 | 0.67 | 0.50 |
| Ramachandran outliers (% ) | 0.00 | 0.00 | 0.00 |
| Rotamer outliers (% ) | 3.48 | 3.39 | 1.88 |
| Clashscore | 2.45 | 4.47 | 3.22 |
| Average B-factor (Å$^2$) | 51.04 | 33.07 | 36.83 |
| macromolecules | 50.74 | 32.10 | 35.78 |
| ligands | 69.77 | 47.86 | 47.82 |
| solvent | 48.45 | 40.91 | 43.15 |
| Number of TLS groups | - | 6 | 8 |

### 7 Bioinformatics

| Sequence name | Comments | Internal/interface interaction? |
|---|---|---|
| $\pi$-$\pi$ interactions | | |
| 1oh8 B446B509 | End of CC region of an antiparallel dimer | yes |
| 1zrt C199P199 | Non-interface (gade) positions | no |
| 2gd7 A39B39 | End of CC region of a parallel dimer | yes |
| 2yiu A199D199 | Non-interface (gade) positions | no |
| 5htf A48B48 | Outside CC area | no |
| 6b02 B289B358 | End of CC region of an antiparallel dimer | yes |
| 6hed L44M44 | Causes kink in parallel dimer | yes |
| 6iko A251B251 | Buried interaction in an antiparallel 6-helix barrel | yes |
| OH-OH interactions | | |
| 1ory A1026A1086 | Buried interaction in an antiparallel 5-helix bundle | yes |
| 2gd7 A39B39 | pi-pi interaction | no |
| 2osz A346B346 | Buried interaction in an antiparallel 4-helix bundle | yes |
| 2qfc A95A119 | Antiparallel dimer | yes |
| 2vzb C33D33 | Non-interface (gade) positions | no |
| 2yfd A88B88 | Non-interface (gade) positions | no |
| 3klt C291D291 | Buried interaction in an antiparallel 4-helix bundle | yes |
| 4aya A43B43 | Non-interface (gade) positions | no |
| 4bt8 B199B230 | Non-interface (gade) positions | no |
| 4h73 E75G75 | Non-interface (gade) positions | no |
| 4h73 F75H75 | Non-interface (gade) positions | no |
| 4jq5 I346J346 | Antiparallel dimer | yes |
| 4ut1 A152A594 | Found in the break between coiled-coil regions | no |
| 5eeb A75C75 | Non-interface (gade) positions | no |
| 5eeb B75D75 | Non-interface (gade) positions | no |
| 5eeb E75G75 | Non-interface (gade) positions | no |
| 5hyl A88C88 | Non-interface (gade) positions | no |
| 5hyl B88D88 | Non-interface (gade) positions | no |
| 6jpf A98A102 | Same helix i+4 interaction | no |
| 6jpf B98B102 | Same helix i+4 interaction | no |

**Table S 7.1** Results from visual inspection of Tyr-Tyr interactions found in coiled coils from the CC+ database. Hits found from $\pi$-$\pi$ distances ≤ 4 Å and hydroxyl-hydroxyl distances ≤ 3.2 Å.

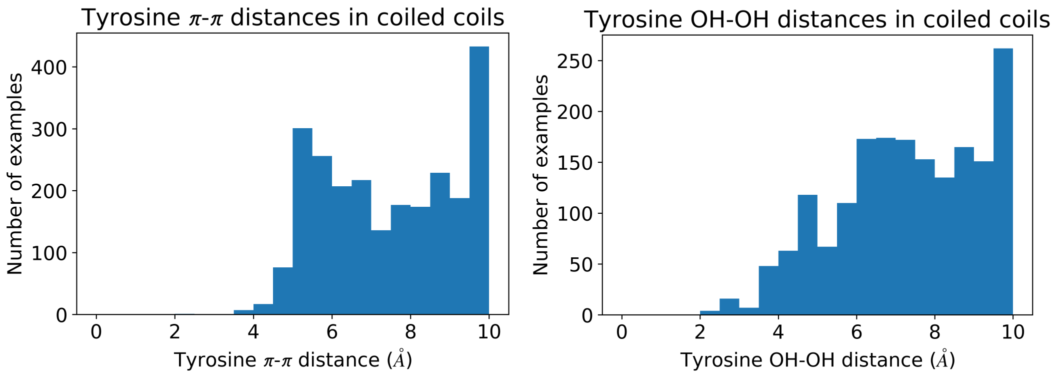


**Figure S 7.1** Histograms displaying Tyr-Tyr distances for $\pi$-$\pi$ and hydroxyl-hydroxyl interaction found in the CC+ database. Data cut at 10 Å distance.

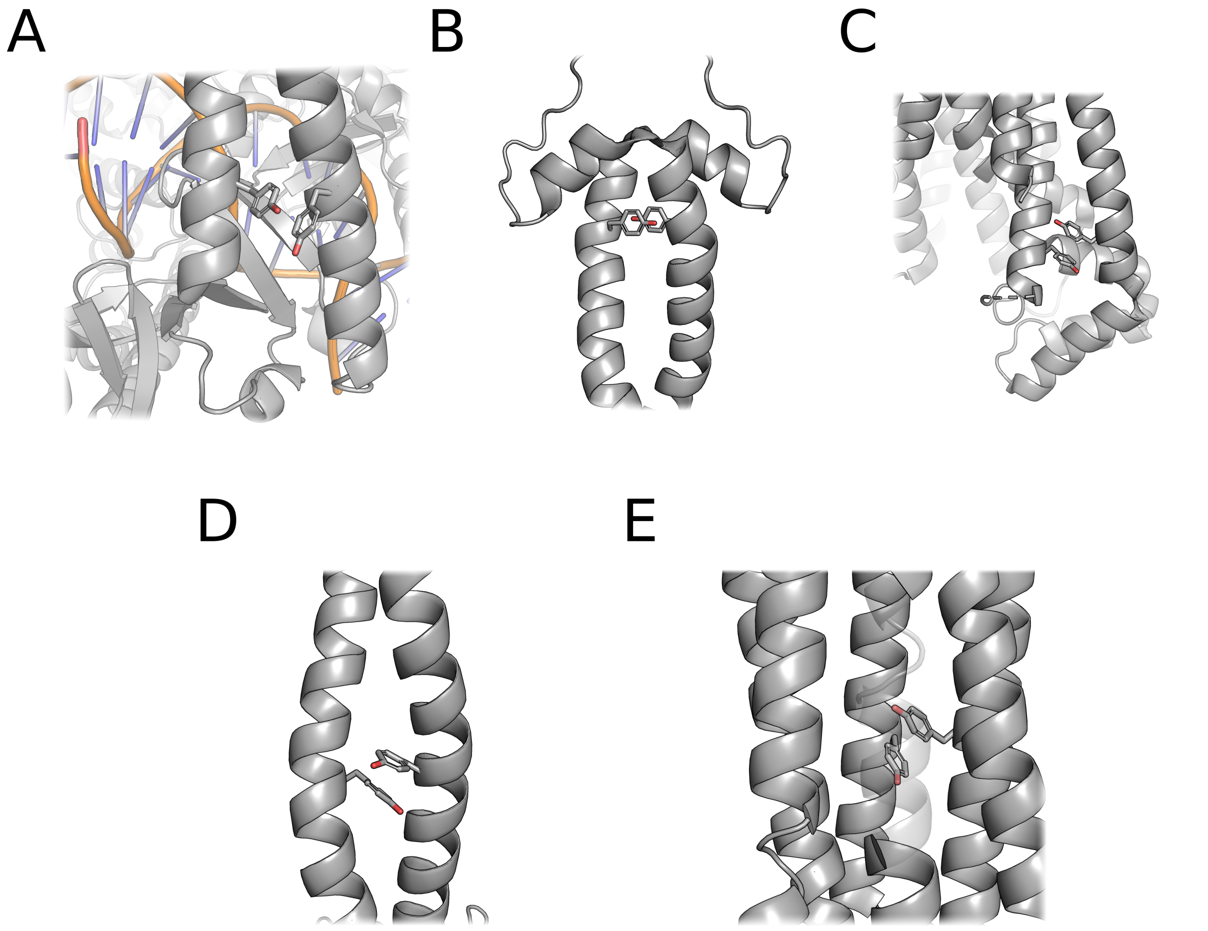


**Figure S 7.2** The five examples of tyrosine $\pi$-$\pi$ interactions found in the CC+ database. Interactions were found in PDB id codes (A) 1oh8 residues B446 & B509, (B) 2gd7 residues A39 & B39, (C) 6b02 residues B289 & B358,(D) 6hed residues L44 & M44, and (E) 6iko residues A251 & B251. Some residues are presented in a semi-transparent form to aid visualisation.

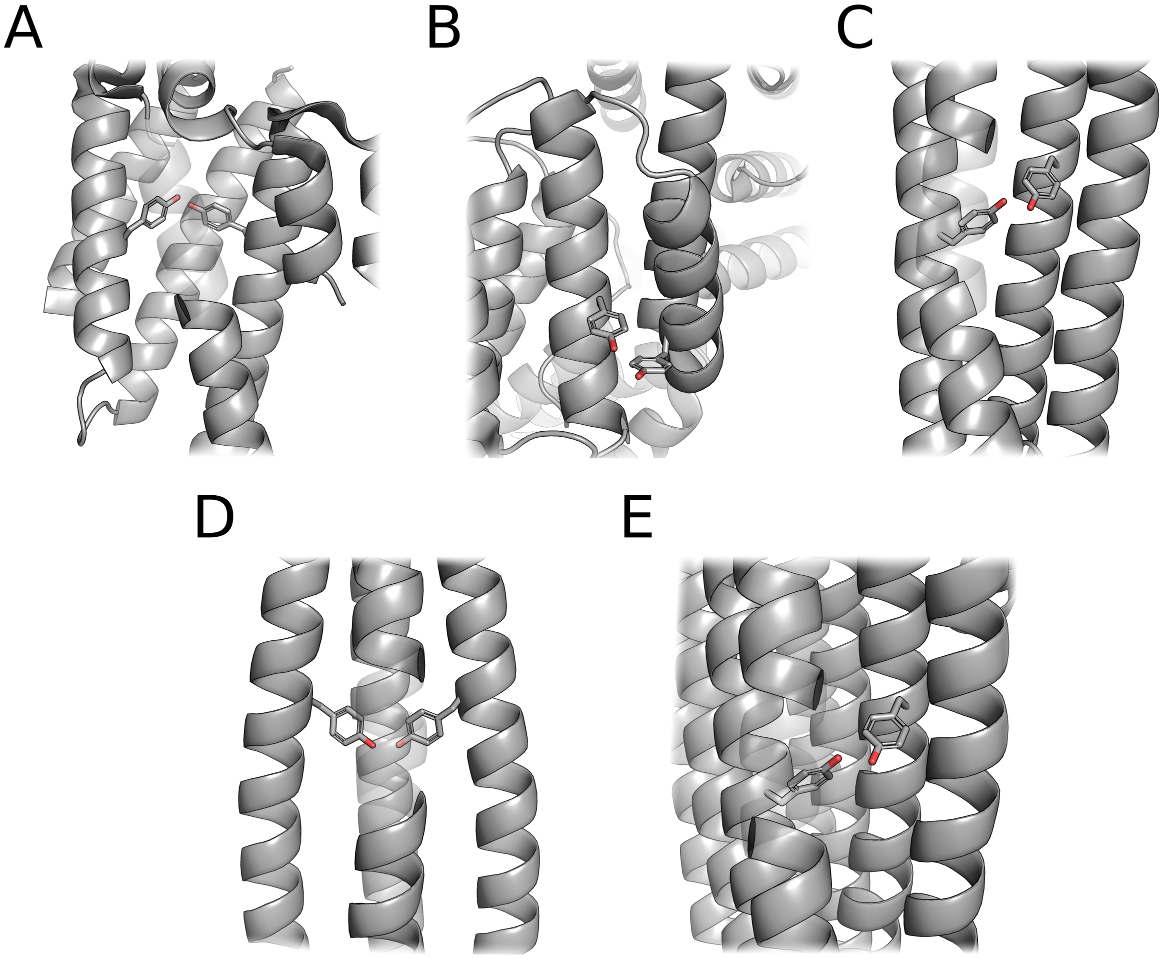


**Figure S 7.3** The five examples of tyrosine hydroxyl-hydroxyl interactions found in the CC+ database. Interactions were found in PDB id codes (A) 1ory residues A1026 & A1086, (B) 2qfc residues A95 & A119, (C) 2osz residues A346 & B346, (D) 3klt residues C291 & D291, and (E) 4jq5 residues I346 & J346. Some residues are presented in a semi-transparent form to aid visualisation.
